# Symmetry breaking of hPSCs in micropattern generates a polarized spinal cord-like organoid (pSCO) with dorsoventral organization

**DOI:** 10.1101/2021.09.18.460734

**Authors:** Kyubin Seo, Subin Cho, Ju-Hyun Lee, June Hoan Kim, Boram Lee, Hwanseok Jang, Youngju Kim, Hyo Min Cho, Sanghyuk Lee, Yongdoo Park, Hee Youn Kim, Taeseob Lee, Woong-Yang Park, Yong Jun Kim, Esther Yang, Dongho Geum, Hyun Kim, Jae Ryun Ryu, Woong Sun

## Abstract

Brain organoid research is advancing, but generation of organoids with proper axis formation, which could lead to spatially ordered structures for complex brain structure and function, still remains a challenge. Axis formation and related spatial cell organization in the CNS are initiated by the symmetry breaking during the early embryo development. It has been demonstrated that the geometrically confined culture of human pluripotent stem cells (hPSCs) can be used to induce symmetry breaking and regionalized cell differentiation. In this study, we generated a polarized spinal cord organoid with a self-organized dorsoventral (DV) organization, using 2D cell patterning by geometric confinement. Initially, the application of caudalization signals to hPSCs promoted the regionalized cell differentiation along the radial axis and sprouting-like protrusion morphogenesis in cell colonies confined to ECM protein micropatterns. Detachment of colonies turned them into extended spinal cord-like organoids which maintained center- and edge-derived two poles. Further analyses including single cell RNA sequencing and spatial transcriptome analysis unveiled that these organoids contained rich repertoire of developing spinal cord cells and exhibited the spatially ordered DV domain formation along the long axis without external organizing signals. Modulation of BMP and Shh signaling can control the extent of DV coverage in organoids following the principles of embryo patterning. Our study provides a simple, and precisely controllable method to generate spatially-ordered organoids for understanding of biological principles of cell patterning and axis formation during neural development.

## INTRODUCTION

Brain organoids are self-assembled three-dimensional cell aggregates derived from pluripotent stem cells (PSCs), which have now become an essential model system for human brain research as they recapitulate the cells and tissue structure of the brain, reflecting developmental trajectory. Much progress has been made in generating organoids of multiple brain regions^1, 2, 3, 4, 5, 6, 7, 8^ with diverse cell types^9, 10, 11^. Bioengineered materials and fusion methods have been applied to develop more complex brain organoids^2, 5, 12, 13, 14, 15, 16, 17, 18^. One of the challenges in current organoid research is the establishment of the anteriorposterior (AP), dorsoventral (DV), and mediolateral (ML) axes, which are crucial in the spatial organization of tissues and organs during embryo development. This fundamental patterning involves a break in the initial molecular and cellular symmetry, leading to precise positioning of signaling centers. Signaling molecules released from these centers are distributed as a gradient, which allows cells to acquire discrete regional identities. Development of AP polarity has been reported in elongated gastruloids and 3D culture systems using blastocysts or epiblasts^19, 20, 21, 22, 23^, and DV patterning is established in brain assembloids^13, 14, 15, 24^. To create a morphogen gradient in a highly controllable way, an engineered signaling center was introduced into one pole of forebrain organoids ^25^ and a microfluidic approach has been utilized^26, 27, 28^.

Symmetry breaking and cell patterning using hPSCs have been relatively well studied in two-dimensional (2D) culture with geometric confinement. Simply confining cells to 2D micropatterns generates a spatially organized signaling environment in a controllable and reproducible manner, leading to regionalized cell fate patterning^29, 30, 31, 32^. The spatial order of BMP4 signaling on micropattern is due to density-dependent receptor relocalization and reaction-diffusion of noggin^33^. Micropattern systems are also useful for studying the dynamics of morphogen signaling events during gastrulation^34^. When micropatterned colonies stimulated with Wnt and activin are grafted into chick embryos, they function as organizers, inducing secondary axis and neural tissue in the host^35^. Additionally, micropattern technologies have been applied to recapitulate early human neurulation^36, 37^. All these studies provide good evidence that pre-patterned geometric confinement can be a useful system for studying symmetry breaking and embryonic patterning using hPSCs.

Recently several groups have reported the generation of spinal cord organoids^1, 2, 6, 7^, but none of them contain spatial patterning established in vivo spinal cord. In the present study, we took advantage of micropattern-based symmetry breaking to generate an organized spinal cord organoid in which DV domains were spatially ordered. The initial cell patterning induced by caudalization signals in 2D geometrically confined colonies was autonomously developed into DV domains in caudalized 3D structures with unique sprouting-like protrusion morphogenesis. The characteristics of spatially organized organoids were analyzed with single-cell RNA sequencing, immunostaining with spinal cord markers, and spatial transcriptomics, and were validated with DV axis perturbation experiments. We suggest that the spatial patterning initially derived from 2D geometrical confinement can be used for the axis formation in polarized 3D organoids, and will be a useful tool for studying early human spinal cord development.

## RESULTS

### Spatial cell patterning in geometrically confined human embryonic stem cell (hESC) colonies

To generate polarized caudal 3D structures, we first induced 2D spatial patterning of caudal neural cells using geometrically confined hESC colonies following caudal neural induction protocols^6, 37, 38^. Microcontact printing was utilized to generate micropatterns of matrigel-coated adhesive circles with varying diameters between 150 and 700 μm on glass coverslips (Fig. 1A). Treatment of micropatterned hESCs with SB431542 (SB) 10 μM and CHIR99021 (Chir) 3 μM, led to a marked protrusion of the central part of the colony, morphologically similar to ‘sprouting up’ in colonies between 200 and 350 μm in diameters (Fig. 1B, S1A and Movie S1). In 500- and 700-μm colonies, instead of center protrusion, multiple low protrusions were seen at the center (Fig. S1B). Colonies of 350 μm in diameters with approximately 50% confluency at the start of neural induction (day 0) exhibited a typical sprouting morphogenesis of approximately 70% yield, and we used mainly 350 μm diameter micropatterns in this study.

**Fig. 1.**
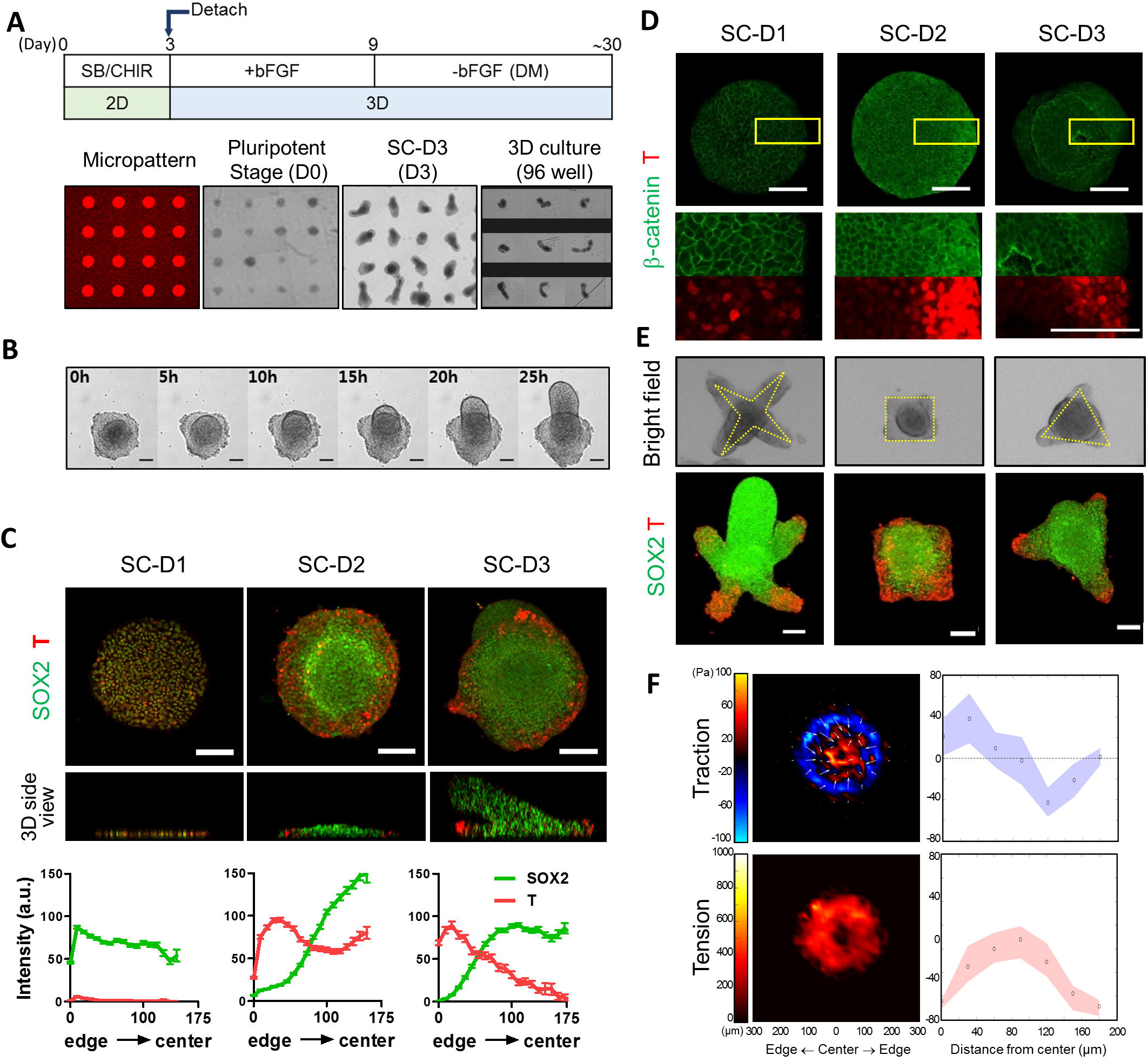
Spatial cell patterning and morphogenesis in geometrically confined hESC colonies. (A) Schematic of the culture protocol. (B) Bright field images of colony morphogenesis developing during the 3rd day of SB/Chir treatment. (C) Immunofluorescence analysis of SOX2 and T expressions. Quantification of fluorescent intensities at each position shown as mean ± SEM. (D) Localization of GFP-β-catenin in micropatterned colonies of β-catenin-EGFP hiPSC cells during SB/Chir treatment. High-magnification images shown at the bottom rows. (E) Effect of micropattern shapes on spatial cell patterning and colony morphogenesis at SC-D3. Yellow dotted lines indicate original micropattern shapes. (F) A color coded map of radial coordinated cellular traction force with traction direction (white arrows) and tension at day1 of SB/Chir treatment and quantification. The mean (black circle) and mid-quartile (blue and red) obtained along the radial lines from the center to the edge of the cell colonies are shown. All images are representative examples from at least three independent experiments. Scale bar, 100 μm.

Next, we examined whether this morphological differentiation of micropatterned colonies was related with spatial patterning of differentiating cells with SB/Chir. hESCs under standard culture conditions were differentiated into neuromesodermal cells characterized by the co-expression of SOX2 and T with SB/Chir treatment (Fig. S2A). At day 2 of SB/Chir treatment (SC-D2), high expression level of SOX2 observed in most of cells in micropatterned colonies at SC-D1 were maintained only at colony centers, while T-positive cells with low SOX2 level progressively emerged at the edge of colonies (Fig. 1C and Movie S2). At that time, initiation of gradual cell accumulation was observed at the colony center, which seemed markedly darker compared with the colony edge (Fig. 1B 0 h and Movie S1). At SC-D3, “center” and “edge” cells were more decisively segregated into two distinct populations. Noticeable protrusions were observed at colony centers, which grew upward over time.

We examined whether spatial cell patterning in a colony results from differential response of cells to an inducer of cell fate. Chir is an activator of Wnt signaling that regulates the stability and nuclear import of β-catenin. While β-catenin was localized at the cell-cell junction in pluripotent ESCs, it was redistributed at SC-D2 into the cytosol and nucleus of cells, notably at the colony edge that expressed high levels of T (Fig. 1D). In SOX2-positive cells at the colony center, β-catenin was localized at the cell-cell junction, suggesting that cellular response to Chir varies from colony center to edge, leading to the induction of different types of cells. The resulting “edge” T-positive cells lost tight junctions (Fig. S2B) and expressed Slug, which is known to regulate the expression of genes responsible for the epithelial-mesenchymal transition (Fig. S2B). On the other hand, SOX2-positive cells of the emergent protrusions at colony centers formed NCAD-positive neural rosettes (Fig. S2C). All these data indicate that spatial differentiation patterning might occur through interplay between self-organizing activity of the hESCs and cues from the geometric boundaries.

This correlation between spatial patterning of SOX2/T expression and emergent center protrusion in micropatterned colonies upon differentiation was shown in colonies exhibiting different morphologies. The SOX2- and T-positive cells were randomly mixed in flat (no protrusion) colonies which were presumably derived from the colonies with low initial cell density, and few T-positive cells existed in dome-shapes (global expansion without center protrusion) colonies (Fig. S1C and Movie S3). All other colonies, on the other hands, with ring- (weak center protrusion) and bell- (typically elongating center protrusion), exhibited spatial patterning of SOX2- and T-positive cells (Fig. S1A, S1C and Movie S4). Especially, colonies from large diameter micropatterns (500-700 um; Fig. S1B) exhibited clear spatial cell patterning with multiple SOX2-positive protrusions. Formation of cell patterning and center protrusion was also achieved in colonies with different micropattern shapes including four-point star, triangle, and square (Fig. 1E). Collectively, these results indicate that geometric confinements faithfully triggered spatial cell patterning, while center protrusion morphogenesis requires emergence of appropriate physical forces triggering the morphogenesis. Accordingly, we found that differential cellular physical force between colony center and edge cells during SB/Chir treatment could contribute to center protrusion. The traction map of the colony showed that cells at the edge developed inward pulling forces, indicating that cells tended to migrate out of the colony. Additionally, tension map showed that high tension developed at the middle of the colony where cells started to aggregate and polarize. Based on these cellular force maps, it was clear that cells at the colony edge and center showed differential force development and distribution (Fig. 1F).

### Development of 3D structures from micropatterned colonies

At SC-D3, we gently lifted differentiated colonies with a center protrusion from the coverslips without disruption of their overall structures using non-enzyme-based depolymerizing solution. Each colony was transferred to ultralow attachment 96 well plate and cultured continuously in the presence of basic fibroblast growth factor (bFGF), which is required for both induction of neural progenitor cells and posteriorization of neuroectoderm, for additional 6 days (bFGFD1–6) (Fig. 2A). The 3D structures underwent gradual reorganization, especially at the bottom part (Movie S5). The entire 3D structure took the shape of peanuts and further extended with a few bendings along their long axes in the presence of bFGF. The top parts (T) with densely packed cells, which had been the protrusion parts in 2D culture, were distinguishable from the bottom parts (B) with a rather brighter appearance, which originated from the disk-shaped bottom.

**Fig. 2.**
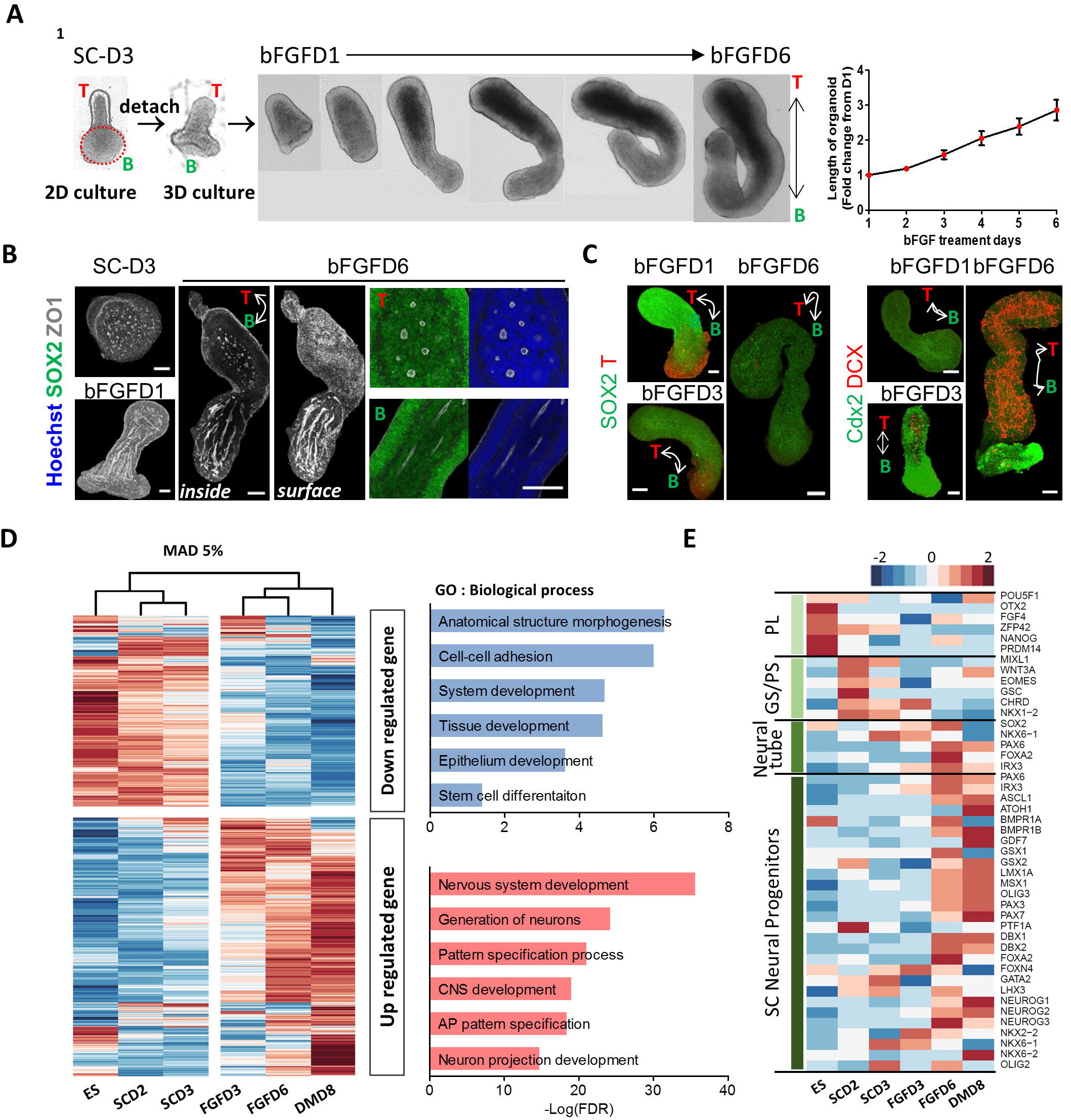
Characterization of polarized cell patterning and temporal gene expression profiles in 3D structures derived from 2D micropatterned colonies. (A) Growth of 3D structures detached from 2D micropatterned colonies in the presence of bFGF for 6 days. Upper protrusive part labeled as Top (T), and the part attached to the micropatterned substrate as Bottom (B). Red dotted circle indicates the initially micropatterned area. Quantification of length of organoids during FGF treatment shown as mean ± SEM. (B) Immunofluorescence analysis of an attached colony (SC-D3) and 3D structures (bFGFD1 and D6). White double arrowhead labeled as T and B indicates the long axis of organoids. Merged images of SOX2/ZO1 and Hoechst/ZO1 with high-resolution show the rosette structures. (C) Immunofluorescence analysis of indicated proteins in the 3D structures at indicated time. (D) Cluster analysis of microarray datasets using time-pooled 2D colonies and 3D structures. Top 5% most variable genes across samples determined by the mean absolute deviation (MAD) function and clustered. The heatmap shows expression profiles of top 5% most variable genes over time with functional annotation. (E) Heatmap of gene expression associated with development over time. Well-known marker genes for pluripotency (PL), gastrulation, and primitive streak (GS/PS), neural tube, and spinal cord (SC) progenitors were examined at each stage. All images are representative examples from at least three independent experiments. Scale bar, 100 μm.

We explored the difference between top and bottom part of the 3D structure. Two distinct types of rosette were developed in each part: The lumens that were marked with ZO1 and surrounded with SOX2-positive cells displayed a circular shape at the top part of a 3D structure, while lumens at the bottom part had a highly extended shape (Fig. 2B and Movie S6). While the entire 3D structure contained SOX2-positive cells, T-positive cells were concentrated at the bottom part and continuously decreased over time (Fig. 2C). The bottom domain containing T-positive cells expressed cdx2, which is a member of the caudal-related homeobox transcription factor ^1, 39, 40, 41^. DCX-positive cells emerged at bFGFD3 and continuously increased (Fig. 2C). This set of observations indicated that cells of micropattern-derived 3D structures were differentiated into neural stem/progenitor cells during bFGF treatment, and the distinct spatial patterning achieved in 2D geometric confinement was further developed into the polarized 3D structures that contained the “top” and “bottom” parts. After 6 days of bFGF treatment, 3D structures were cultured without bFGF for further differentiation into neurons. We referred to this stage of growing without bFGF as the DM (differentiation medium) stage.

To assess the differentiation direction and potential of 3D structure development, microarray analysis was carried out using samples collected at varying culture stages: the pluripotent stage (ES), days 2 and 3 of SB/Chir treatment (SCD2 and SCD3), days 3 and 6 of bFGF treatment (FGFD3 and FGFD6), and day 8 of differentiation medium stage (DMD8) (Fig. S3A). Genes associated with central nervous system development, AP pattern specification, and neuron-related process were highly transcribed in FGFD3, FGFD6, and DMD8 samples. Genes downregulated over time were involved in cell-cell adhesion, tube development, and epithelium development (Fig. 2D). Specifically, genes associated with gastrulation and primitive streak were initially expressed, and then neural tube-associated genes increased (Fig. 2E). Marker genes for neural progenitors in the spinal cord started expressing at FGFD6. HOX gene clusters, up to HOX10, were gradually expressed, suggesting that our 3D structures held the caudal identity beginning from the hindbrain to the lumbar vertebra (Fig. S3B). Pathway and ontology analyses with Enrichr^42, 43, 44^, an open bioinformatics platform that utilizes mammalian data, showed that spinal cord-related genes were enriched in the top 250 most variable gene set, and interestingly, neural crest (NC)-related genes were induced by our protocol (Fig. S3C). NC cells arise from the neural plate border, and after neural tube closure, they delaminate from the region between the dorsal neural tube and overlying ectoderm in vivo during development. Therefore, it appears that the NC cells were induced by our protocol aimed at directing caudal neural fate^39, 45^. Taken together, these microarray analysis data indicated that our stepwise induction and differentiation protocol produced early caudal neural stem/progenitor cells through the developmental stages of gastrulation and neurulation, which possibly matured into spinal cord cells. Spatial patterning established in 2D patterned substrates developed into a polarized 3D structure composed of top and bottom parts. Therefore, we termed our 3D structures “polarized spinal cord organoids (pSCOs)” that were reminiscent of the early developmental stage of the embryonic spinal cord.

### Single-cell RNA-sequencing (scRNA-seq)

To assess the full variety of cell types present in the pSCOs and their temporal changes in an unbiased manner, scRNA-seq was performed using 10,618, 11,690 and 13,696 cells in the FGFD1, FGFD6 and DMD10 samples, respectively. We used a nonlinear dimensionality reduction visualization algorithm, uniform manifold approximation and projection (UMAP), to visualize cells and determine their identity. Cell type annotation was performed following the diagram of spinal cord development by Alaynick *et al*.^46^. All single cells from 3 stages grouped into six main clusters: 1) Neural progenitor, 2) mitotic dorsal, 3) mitotic ventral, 4) postmitotic dorsal, 5) postmitotic ventral, and 6) mesoderm-like clusters (Fig. 3A and 3B). Top 10 differentially expressed genes per cluster were shown in Supplementary data 1. A higher percentage of postmitotic SC neurons existed in the DMD10 sample, while the FGFD1 sample and FGFD6 samples showed a higher percentage of neural progenitors and mitotic SC progenitors, respectively (Fig. 3A and Fig. S4A), indicating that organoids were developing progressively over time. Further sub-clustering of each cluster showed that the organoids contained almost the full repertoire of spinal cord domains at both mitotic and postmitotic stages (Fig. 3C, Fig. S4B and Supplementary data 2). In the mitotic dorsal cluster, cells of the roof plate and dorsal domains from pd1 to pd6 were identified, and ventral domains from p0 to p2 and pMN were identified in the mitotic ventral clusters. dI1, dI2, and dI3 domains were identified in the postmitotic dorsal clusters and ventral domains V0, V1, V2, and MN were identified in the postmitotic ventral clusters. In addition, a gradual increase of HOX gene expression over time confirmed the caudal identity of our organoids (Fig. S4C). For further validation of the cluster assignment, previously annotated clusters from the developing mouse spinal cord^47^ was compared to our pSCOs (Fig. S5A and S5B). Overall, cluster annotation of mitotic and postmitotic cells in our pSCOs matched the spinal cord clusters.

**Fig. 3.**
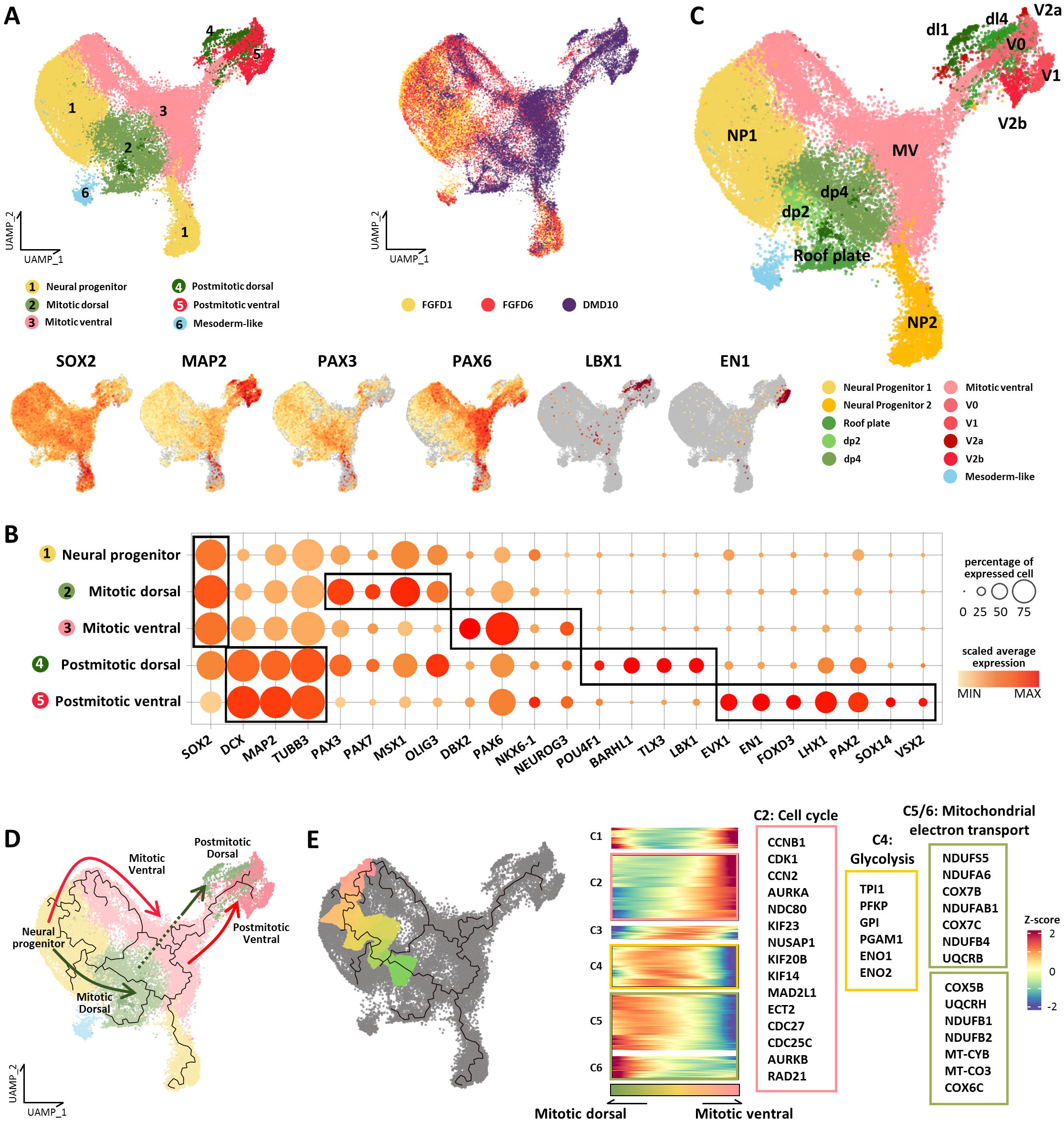
Molecular characterization of the organoid cell population via single-cell RNA-sequencing (scRNA-seq) (A) UMAP plot of integrated FGFD1, FGFD6 and DMD10 datasets. The six main cell clusters (left UMAP plot) and cells of different stages (right UMAP plot) are differently colored. UMAP plots at the bottom row show gene expression levels of representative signature genes for each cluster. (B) Dot plot showing the expression of representative signature genes across five main clusters. The size of each circle indicates the percentage of cells in each cluster where the indicated gene was detected, and the color intensity reflects the scaled average expression level within each cluster. Genes in black boxes are representative signature genes in each cluster. (C) Sub-clustering of six main clusters in Fig. 3A. Mitotic and postmitotic cells of organoids were differently colored according to expression profiles of dorsal and ventral spinal cord-related genes. Domain-unspecified cells were labeled with mitotic ventral. (D) Monocle-based trajectory analysis of pSCOs. Each trajectory path are differently colored. (E) Dorsal –Ventral split in the neural progenitor cluster. Pseudotemporal gene expression profiles were hierarchically clustered. Genes related with indicated GO terms in each cluster are highlighted at right side.

Monocle-based trajectory analysis was performed to further assess the developmental progression of cells in organoids and we constructed a differentiation trajectory from the scRNA-seq data^48^. This analysis revealed the trajectory starting from common neural progenitors to dorsal and ventral SC neurons through the fate commitment toward dorsal and ventral progenitors (Fig 3D). To investigate the process of initial dorsal and ventral cell fate commitment, we examined pseudotemporal transcriptome profiles associated with alternative cell fates, focusing the bifurcation separating dorsal and ventral branches at neural progenitor clusters (Figure 3E). Differential expression patterns along the trajectory were grouped into 6 clusters. Alignment of them with directions toward dorsal and ventral progenitors in pseudotime made us to speculate primitive neural progenitors (cluster 4, C4), and ventrally (C2-3) or dorsally (C5-6) specifying progenitors. Their DEG analyses revealed that genes associated with canonical glycolysis process were enriched in the primitive progenitors (C4), and genes related with mitochondrial electron transport were enriched in the clusters specifying toward dorsal progenitors (C5-6). This indicates that energy shift from glycolysis to mitochondrial oxidative phosphorylation gradually occurs during the transition from neural progenitors to dorsal progenitors^49, 50^. On the other hand, genes associated with cell cycle and mitosis were enriched in neural progenitors committed toward ventral fate (C2) (Supplementary data 3)^51^.

### Spatial transcriptomics (ST) of pSCOs

The polarized structure observed in early 3D structure development (Fig. 2B) led us to hypothesize that ventral and dorsal cells identified by the scRNA-seq analysis were segregated spatially in the organoids. VISIUM spatial transcriptomics (ST) was applied to explore spatial characterization of the organoids in an unbiased manner using DMD10 sample (Fig. 4A). Specifically, the spatial pattern of gene expression within a 10 μm-thickness section of an organoid along its long axis at a resolution of 53 spots was analyzed. One spot with a 55-μm diameter contained approximately 15 cells. The application of UMAP to all 53 spots across the section resulted in the identification of two main clusters. Spots in one cluster that covered the bottom half of the section were expressing genes related to SC dorsal domains, while those in the other cluster that were located at the top half of the section were expressing genes of SC ventral domains. Next, the ST data was integrated with the scRNA-seq data using an “anchor”-based integration method that enables the probabilistic transfer of annotations from the scRNA-seq data to the ST data^52^. This returns a prediction score for each cell type for every spot in the ST dataset that ranges from 0 to 1. Each spot in the ST data could be considered a weighted mix of cell types identified by scRNA-seq. Using prediction scores, we found that dorsal spinal cord cells identified by the scRNA-seq analysis were located mainly at the bottom side of the organoid section (Fig. 4A). On the other hand, ventral domain cells identified via the scRNA-seq analysis were mostly located at the top side, indicating that our organoids were anatomically distinguished by different transcriptomes. Subsequently, we used marker genes and differential gene expression analysis to spatially assign cell types. Among mitotic spots with high levels of SOX2, spots highly expressing mitotic dorsal markers (PAX3, PAX7, ASCL1, and GSX2) were located at the bottom side (dorsal side) and labeled together as pd3, 4, and 5 domains (Fig. 4B). dI1, 2, and 3 domains highly expressing POU4F1, LMX1B, and dI4, 5, and 6 domains expressing LBX1, PTF1A, and GSX2 were identified at the spots located at the bottom side (Fig. 4C). In contrast, spots expressing PAX6 and DBX2 at the top side were labeled as the p0 domain, while Nkx6-2 (p1), Nkx6-1 (p2), and Neurog3 (p3) were expressed at the top spots (Fig. 4B). The top spots also contained postmitotic ventral domains expressing marker genes such as V0 (EVX1), V1 (EN1), V2a (VSX2, SOX14, and SOX21), V2b (GATA2), and V2c (SOX1) (Fig. 4C). In addition, Mnx1 was highly expressed in the MN domain at the top spots. Collectively, this spatial transcriptome analysis revealed that our pSCOs have dorsoventral patterning, but partially. On the other hand, spatial expression of HOX genes demonstrated that posterior HOX genes were strongly expressed in the entire organoid section (Fig S6A), indicating that our organoids are broadly specified to spinal cord regions^53, 54^. However, we also discovered that the ‘bottom’ part of the pSCO which contained dorsal-like cells exhibited relatively stronger HOX9 expression, which is in consistent with our finding that stronger HOX expression was a signature of dorsally confined neural progenitors.

**Fig. 4.**
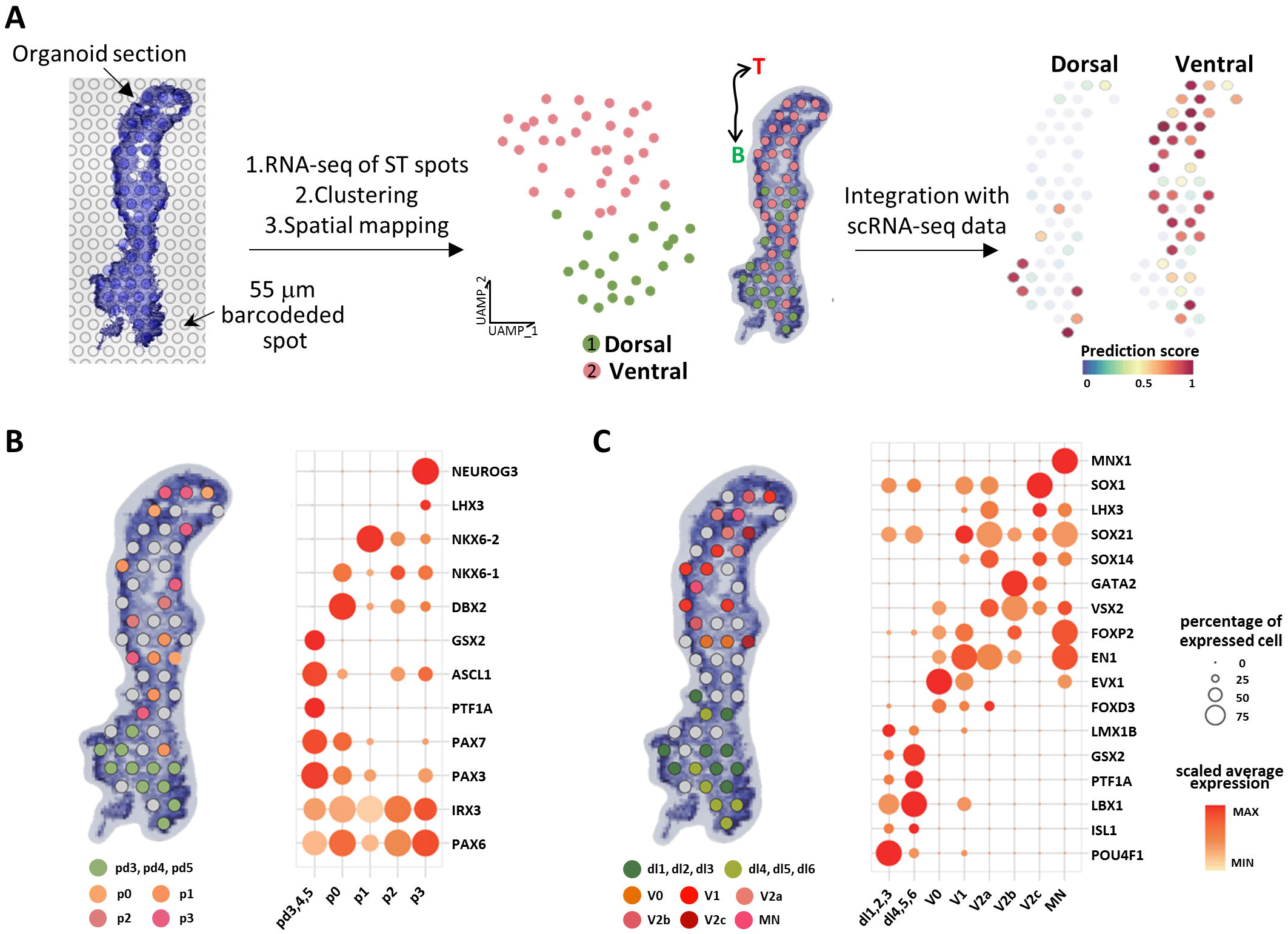
Molecular characterization of the organoid cell population via Spatial transcriptomics (ST) (A) Spatial transcriptomic analysis of the DMD10 organoid. Frozen 10 μm sections of DMD10 organoid are placed onto a slide with barcoded spots, and sequencing, clustering and mapping of 53 spots in organoid section are done. Two main clusters are mapped with different colors. Dorsal and ventral diagram on the right side shows the spot assignment by prediction score analysis using integrated scRNA-seq and ST datasets. Black double arrowhead labeled as T and B indicates the long axis of organoid section. Mapping of spinal cord mitotic domains (B) and postmitotic domains (C) in the organoid section. Spots are differently colored according to their domain assignment, and unspecified spots are colored in gray. Dot plot showing the expression of representative signature genes across assigned domains.

### Mapping of dorsal and ventral spinal cord domains in the organoids using immunostaining

Next, we obtained a spatiotemporal overview of the cellular organization in the organoids by immunostaining with antibodies against dorsal and ventral markers both at mitotic (Fig. 5A) and postmitotic (Fig. 5B) stages. The regional cytoarchitecture differs between top and bottom parts in the organoids (Fig. 5A ZO1 and Movie S7), which indicates the existence of different cell populations. The fact that these rosette-forming cells in the entire organoid were all SOX2-positive neural progenitor cells excluded the possibility that different stages of cells were populated at the top and bottom parts of our organoids. Dorsal domain markers such as Pax3, Pax7, and Olig3 displayed polarized regionalization at the bottom part of the organoids (Fig. 5A and Movie S7). In a later stage, this bottom part expressed high levels of Bran3a, a marker for dorsal neurons (dI1-3) (Fig. 5B and Movie S8). In contrast, Nkx6-1- and Pax6-positive cells were preferentially distributed at the top (Fig. 5A and Movie S7), where postmitotic ventral markers including Evx1 (Vo), Chx10 (V2a), or Chat (MN) were regionally expressed (Fig. 5B and Movie S8). These results indicated that the bottom part of our organoids developed from mitotic dorsal progenitors to postmitotic dorsal neurons, and the top part was differentiating into ventral domains although we could not observe the neural tube-like structure, a tubular structure with a central lumen^24^. The existence of NC cells was predicted using our microarray data (Fig. S3C). Indeed, Slug- and/or SOX10-positive NC progenitor cells were present at the bottom (dorsal) part of the organoids (Fig. 5A and Movie S7), and these cells were differentiating into Bran3a/Islet-double positive cells at the bottom (Fig. 5B and Movie S8). In addition, Lhx1/2, which has been known to be expressed both in dorsal and ventral domains in the spinal cord, was expressed in the entire organoids. Further, peripherin-positive neurons were observed in the whole organoids; however, a significant amount of these cells was observed in the ventral part, indicating that these neurons might be motor neurons. Together, the top and bottom parts that were initially distinguished by different cytoarchitectures were spatially patterned into ventral and dorsal domains, respectively, with maturation into spinal cord neurons similar to an embryo spinal cord development (Fig. 5A and 5B diagrams).

**Fig. 5.**
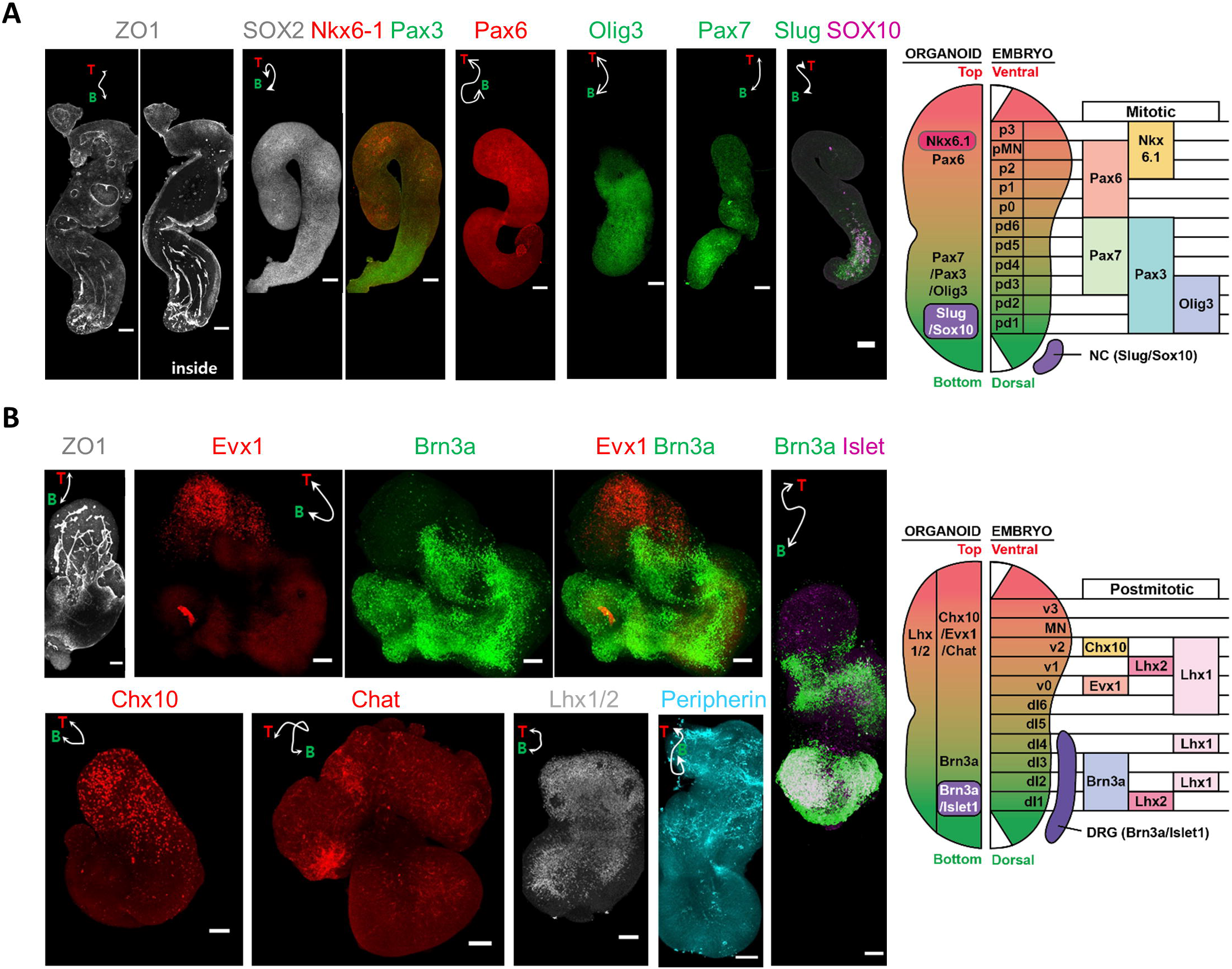
Characterization of spatial dorsoventral patterning of the spinal cord organoid. Immunofluorescence analysis of spinal cord organoids with markers for spinal cord progenitor cells (A) and postmitotic neurons (B). White double arrowhead labeled as T and B indicates the long axis of organoids. Diagram on the right side shows that genes expressed in top and bottom regions in organoids match mitotic ventral and dorsal markers, respectively, in embryonic spinal cord during development. All images are representative examples from at least three independent experiments. Scale bar, 100 μm.

### Regulation of DV axis organization via changes in BMP and Shh signaling activities

The high degree of organization in DV gene expression in our organoids prompted us to examine the distribution of patterning cues in the organoids. The developing vertebrate neural tube is precisely patterned by antiparallel signaling gradients of BMP and Sonic hedgehog (SHH) along the DV axis^46, 55, 56^. In our pSCOs, BMP4 and BMP7 were highly expressed in cells of the mitotic dorsal cluster and dorsal spots, while cells with high expression levels of SHH, although the population was small, were found in the postmitotic ventral clusters and a ventral spot (Fig. 6A). Therefore, we examined whether the disruption of BMP signaling pathway could affect DV organization established in the organoid. Polarized expression of Pax3 in the dorsal part was significantly decreased upon treatment with LDN, a BMP pathway inhibitor, during bFGF treatment (Fig. 6B). BMP inhibition also interfered with organoid growth (Fig. S7A). Perturbed BMP signaling also affected cell specification in the ventral domain. Nkx6.1 expression in the ventral part was significantly increased by the LDN treatment (Fig. 6B), and Nkx6.1-positive cells were found even in the dorsal end of the organoid (Fig. S7B). In contrast, when BMP2 was added to the culture medium, which led to an entire organoid exposed to BMP2, the Pax3-positive domain was expanded to the whole organoid, and the Nkx6.1-positive domain disappeared (Fig. 6B). A SHH inhibitor, GDC 0449 (GDC), inhibited Nkx6.1 expression and enlarged the PAX3-positive domain (Fig. 6B). Activation of the Shh signaling pathway with purmophamine (PMP), a Smo receptor agonist, significantly decreased the size of the Pax3-positive domain and instead expanded the Nkx6.1-positive domains (Fig. 6B) with high expression levels of Nkx6.1. Both BMP2 and PMP treatments promoted organoid growth (Fig. S7A).

**Fig. 6.**
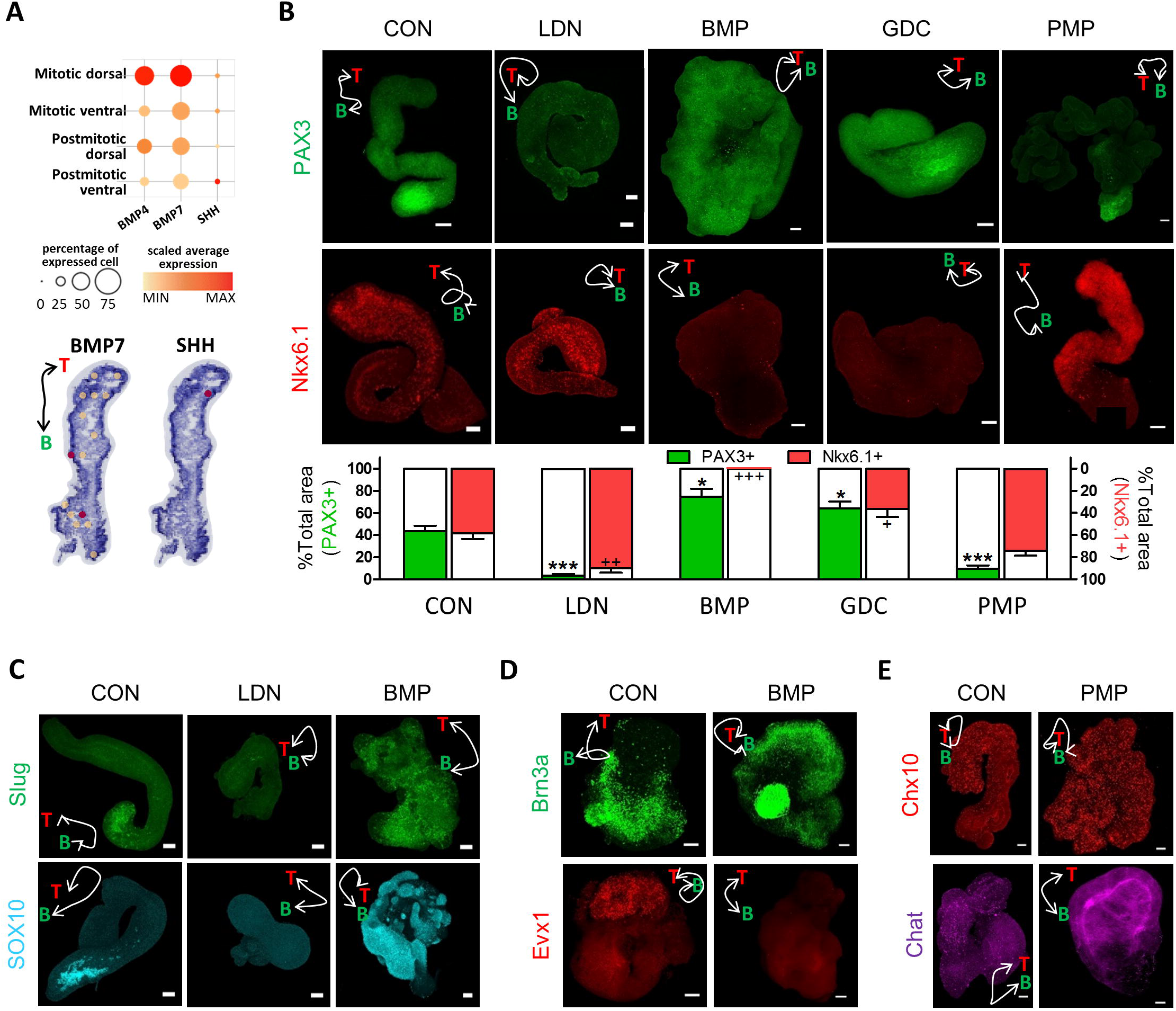
Regulation of dorsoventral (DV) axis organization in the organoids via changes in BMP and Shh signaling activities. (A) Dot plot showing the expression of BMP and SHH across four main clusters and mapping of BMP and SHH in the organoid section. (B) Effect of BMP signaling inhibitor (LDN), BMP, Shh inhibitor (GDC), and Shh activator (PMP) on DV-like axis organization of spinal cord organoids in the progenitor stage. Organoids treated with the indicated reagents were immunostained with the dorsal marker (PAX3) and ventral marker (Nkx6.1). Quantification of PAX3- and Nkx6.1-positive area (% total area) at each condition shown as mean ± SEM. Scale bar, 100 μm. *, +: p<0.05, ++: p<0.01, ***, +++: p<0.001 (C) Effect of BMP signaling inhibitor (LDN) and BMP on the generation of neural crest cells in spinal cord organoids. (D) Effect of BMP on the DV axis organization of spinal cord organoids in the postmitotic stage. Organoids treated with the indicated reagents were immunostained with the dorsal marker (Brn3a) and ventral marker (Evx1). (E) Effect of Shh activator (PMP) on the DV-like axis organization of spinal cord organoids in the postmitotic stage. Organoids treated with the indicated reagents were immunostained with the ventral markers (Chx10 and Chat). White double arrowhead labeled as T (top side) and B (bottom side) indicates the long axis of organoids. All images are representative examples from at least three independent experiments. Scale bar, 100 μm.

BMP signaling is essential for NC development. As expected, NC precursor cells completely disappeared via the inhibition of the BMP signaling pathway; however, they significantly increased upon BMP treatment with uniform distribution along the whole organoid (Fig. 6C and Fig. S7C). It seemed that bFGF signaling is required for the proper development of NC precursor cells (Fig. S7D). Optimal duration of the bFGF treatment was also necessary for proper development of the DV axis in the organoid (Fig. S7E).

We examined the effect of the disturbance of BMP and Shh signaling pathways on DV patterning in the postmitotic neuron stage. Upon BMP treatment, Brn3a-positive dorsal interneuron domains expanded to the whole organoid, while Evx1-positive V0 domains disappeared (Fig. 6D). Upon PMP treatment, the Chx10-positive V2a domains significantly expanded (Fig. 6E). When organoids were treated with PMP during bFGF treatment and differentiated further, Chat-positive motor neurons significantly increased in the ventral part (Fig. 6E). Moreover, Islet 1/2-positive MN domains expanded to more dorsal parts with an increased duration of PMP treatment (Fig. S7F). Taken together, these data indicated that DV patterning in our organoids took place by the interactive regulation of the BMP and Shh signaling pathways similar to those observed in vivo. As fates of dorsal and ventral cells were not completely specified in the bFGF treatment stage in the organoid, fates of these progenitor cells might be interchangeable according to the balance between BMP and Shh signaling pathways regulated by external cues.

## DISCUSSION

Our pSCOs provide new insights into the establishment of spatial patterning in vitro, especially how 2D symmetry breaking develops into 3D polarization, leading to precise positioning of cells in a highly reproducible way. Dynamic processes of development, such as a process of cell specification in early patterning and its maturation to counterpart neurons, could be modeled in organoids in a controllable manner.

Several groups have reported 3D culture systems including gastruloids and organoids that recapitulate symmetry breaking and the characteristic in vivo axis. Autonomous symmetry breaking occurs after bath application of patterning factors to human or mouse PSCs without asymmetry in the initial signal, extra-embryonic tissues, or localized signaling centers^19, 21, 22^, ^23^. On the other hand, to establish an axis in the neural organoids, either addition of signaling factors to the media or an fusion of a signaling-releasing cells with forebrain organoids are necessary^13, 14, 15, 24, 25^. Compared with the above cell patterning strategies, a key feature of our strategy was that we utilized a very simple, easy, and reproducible micropattern technique for initial 2D cell patterning, and we did not include signaling factors to induce DV axis. Spinal cord organoids with patterning have been also previously reported^7, 57^. However, bath application of BMP, Shh or related inhibitors in their studies could not induce dorsal and ventral parts simultaneously within an organoid, as in our study.

Geometric confinement by micropattern is sufficient to induce spatial organization that developed into well-ordered spatial positional domains in our pSCO. Spatial patterning on micropattern has been explained using the ‘reaction-diffusion’ and ‘edge-sensing’ concepts ^33, 58^. Since the relocalization of β-catenin in response to SB/Chir was different at the edge and center cells and T-positive cells were generated at the edge of colonies irrespective of colony size, spatial patterning of SOX2- and T-positive cells in our system might be better explained by the ‘edge-sensing’ model. The difference between edge and center cells in the colony was also demonstrated in terms of physical force. The force map indicated that the combination of the inward pulling force in edge cells and high tension at the colony center might develop the force for cells to sprout up. The spatial patterning of SOX2- and T-positive cells on the micropattern autonomously induced prominent 3D structures, which were detached and subsequently grown in 3D culture. The fact that the polarized structure was established (Fig. 3) gave rise to the speculation that different environments for the emergence of different types of cells were created at the top and bottom regions in the organoids. In fact, BMP and Shh, which are DV patterning factors, were expressed in dorsal and ventral clusters, respectively, as shown in the scRNA-seq analysis and spatial transcriptomics (Fig. 6A). Two regions in the organoid were also distinguished by the different rosette structures and emergence times of DCX-positive cells (Fig. 2B and 2C). Rosette structures have been reported to differ according to the distance from the Shh source in forebrain organoids^25^, which is in consistent with our current observations.

As shown in Fig. 7, our organoids recapitulated neural tube development in vivo in many aspects. Neuroectoderm or neuroepithelial cells that cover the surface of 3D structures and form neural rosette inside organoids sprout up and become ventral parts like ventral part formation from the neural plate at the neural tube fold stage (second row in Fig. 7). After neural tube closure, the dorsal neural tube is isolated from the non-neural ectoderm in vivo. Similarly, the bottom part of the 3D structures detached from the culture plate becomes dorsal domains in organoids (third row in Fig. 7). As NC cells delaminate from the region between the dorsal neural tube and overlying ectoderm in vivo, they are generated in the dorsal-most part of the organoid (four rows in Fig. 7). Finally, neural stem cells at the top part of the organoids become ventral progenitor cells, and cells at the bottom become dorsal cells, leading to dorsal and ventral postmitotic neurons with spatial DV organization reminiscent of the spinal cord DV axis in vivo (bottom row in Fig. 7). Additionally, NC cells differentiate into dorsal root ganglion neurons in vivo. Brn3a/Islet-positive sensory neurons were also observed in the dorsal part of the organoids.

**Fig. 7.**
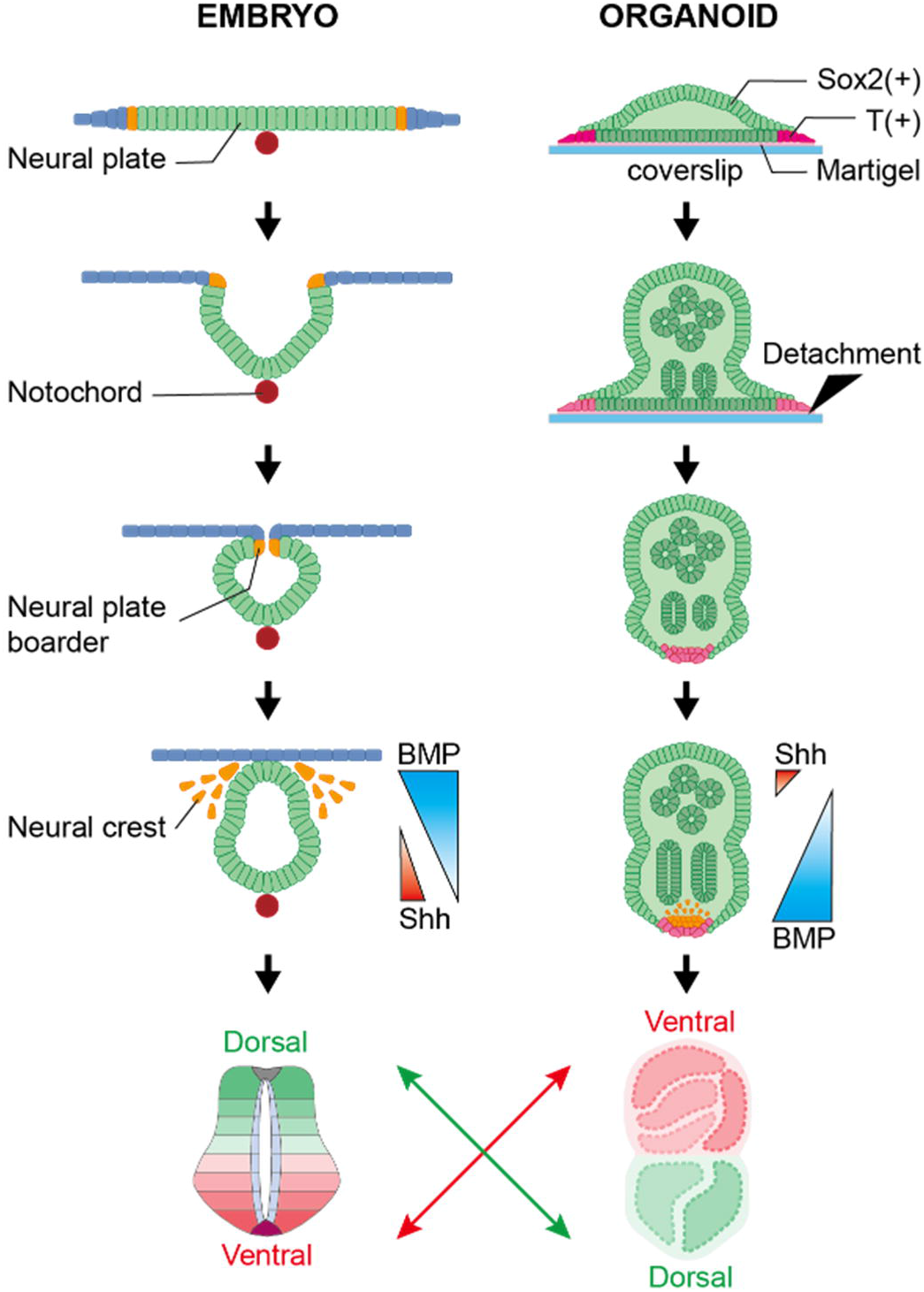
Schematic comparisons of spinal cord DV axis patterning in embryos and polarized DV patterning in organoids. pSCOs recapitulate neural tube development in vivo in many aspects. Neural stem cells at the top part of the organoids become ventral progenitor cells, and cells at the bottom become dorsal cells, leading to dorsal and ventral postmitotic neurons with spatial DV organization reminiscent of the spinal cord DV axis in vivo.

It is yet curious why axial elongation morphogenesis is associated with DV axis. Several reports with gastruloid models suggested that the axial elongation was associated with AP axis, which was evidenced by the gradual patterning of HOX codes. On the other hand, although we observed the quantitative differences in the HOX gene expression levels along the long axis, their difference was rather associated with the stronger HOX expression in dorsal cell populations, and we favor the idea that axial elongation was more strongly associated with DV axis in our study. This phenomenon in our model may be associated with different timing of NMP-derived neural progenitors along the dorsoventral axis^59^, and ‘oblique’ formation of junctional neurulation in caudal spinal cord^60^. Further studies with pSCO model and comparison with human spinal cord development will clarify this issue.

Our strategy to make DV patterning in organoids, however, had a few limitations and remaining issues that need to be addressed in future studies. While we found a rich repertoire of spinal cord cells existed and was spatially organized, we could not observe signaling gradient-dependent organization of dorsoventral domains, as shown in embryonic spinal cord development. In addition, current version of our protocol does not allow the maintenance of the elongated morphology of pSCOs in the extended culture experiments; Often, pSCOs in later stages bend or one part adheres to other parts within or among organoids, which may cause unexpected cell migration or cell mixing over time, leading to the disturbance of the DV axis.

## MATERIALS AND METHODS

### Cell culture and differentiation

For maintenance, hESCs (H9) were grown in the E8 or mTeSR1 (STEMCELL Technologies) medium in tissue culture dishes coated with Matrigel (Corning, 354277; 1:25 in DMEM/F12). Cells were passaged using ReLeSR (STEMCELL Technologies) every 5 days. For experiments, hESCs were grown on coverslips in the mTeSR1 medium 1 or 2 days after passaging. The medium was replaced with the neural cell induction medium (DMEM/F12, 1% N2 supplement, 2% B27 supplement, 1% NEAA, 1% penicillin, and 0.1% 2-mercaptoethanol) containing 10 µM SB431542 (R&D) and 3 µM CHIR99021 (Sigma), and was subsequently replaced every day. After 3 days, they were further grown in 3D with the neural cell induction medium containing 20 ng/mL bFGF. After 6 days of bFGF treatment, 3D structures were grown without bFGF in the neural cell induction medium, which we called the differentiation medium (DM) stage. All informative list of the reagents including antibodies was shown in Table S1.

### Stamp fabrication and micro-contact printing

A stamp for printing biomolecules was fabricated using soft lithography techniques^61, 62^. The photoresist (SU-8 2010, Microchem, USA) was spin coated for 30 s at 1,000 rpm (target thickness: 20 μm) and baked for 4 min at 95°C. The photoresist layer was exposed to UV light using a pre-designed photomask (exposure dose: 175 mJ/cm^2^). After exposure, the wafer was baked for 5 min at 95°C. The wafer was then immersed in an SU-8 developer to remove the uncrosslinked photoresist layer. After the developmental process, the wafer was sequentially rinsed with isopropanol and distilled water. To cast the stamp, the polydimethoxysilane (PDMS) prepolymer:curing agent mixture (10:1 [w/w], Sylgard 184 Silicone Elastomer Kit, Dow Corning, USA) was cured overnight on the SU-8 mold at 60°C. Finally, the cured PDMS was cut into an appropriate size to obtain the finished PDMS stamps. Microstamps with the desired dot pattern were inked with Matrigel (Corning, 354277) and then applied to the coverslips for printing^63^.

### Immunostainings

Cells grown on coverslips were fixed using 4% paraformaldehyde in 0.1 M phosphate-buffered saline (PBS; pH 7.4) for 20 min at 25°C. Cells were washed three times with PBS and then blocked with 0.2% Triton X-100 with 3% bovine serum albumin (BSA) in PBS for 30 min at 25°C. Subsequently, cells were incubated with primary antibodies in a blocking buffer overnight at 4°C. After incubation, the cells were washed three times with PBST (0.2% Triton X-100 in PBS) and afterward incubated with secondary antibodies, with Hoechst for cell nuclei staining (1:2,000), in the blocking buffer for 1 h at 25°C. After washing with PBST, the cells were imaged using a confocal microscope (Leica TCS SP8 confocal microscope).

3D structures were fixed using 4% paraformaldehyde in 0.1 M PBS (pH 7.4) for 1 h at 25°C, washed three times with PBS, blocked with 0.1% Triton X-100 with 6% BSA in PBS overnight at 25°C, and then incubated with primary antibodies in the blocking buffer overnight (twice) at 25°C. After incubation, the cells were washed three times with PBST (0.1% Triton X-100 in PBS) and subsequently incubated with secondary antibodies with Hoechst (1:2000) in the blocking buffer overnight (twice) at 25°C. After washing with PBST, organoids imaged using a confocal microscope (Leica TCS SP8 confocal microscope).

### Time-lapse imaging

Morphogenesis of 3D structures was recorded using the JuLI^TM^ stage (NanoEnTek Corp.), a real-time cell history recorder. Cell images were acquired directly from the culture plate (12 wells) in an incubator automatically at a bright field. A 4× objective lens was used, and the focus was sustained.

### Image analysis and Statistical analysis

Every experiment was repeated at least three times and representative images were shown in Figures. Image analysis was performed with Custom MATLAB (R2017a, MathWorks) and ImageJ. Chart drawing and statistical analyses were performed using the SigmaPlot software and Graphpad Prism.

### Quantitative analysis of kinematics and mechanical dynamics of hESC colony

The movement and physical force of the cell colonies were measured as previously reported (Jang et al., 2017). Briefly, hESCs were patterned on polyacrylamide (Bio-Rad, USA) substrates embedded with fluorescent beads (diameter = 500 nm; FluoSpheres; Invitrogen, USA) in the same manner as described above. Using a JuLI stage live cell imaging system (NanoEnTek, Korea) with a 4× objective lens (Olympus, Japan) in a cell culture incubator, a bright-field channel for the patterned cell colony images and a red fluorescent protein (RFP) channel for bead images were simultaneously collected while culturing the hESC colony. The obtained images were numerically converted and calculated using a customized source code developed with MATLAB (MathWorks Inc., USA). To calculate the cell and bead displacements, particle image velocimetry analysis was conducted on each image set. The displacement result from the bright field images was converted to a movement trajectory and velocity of the cells within the colony, and the displacement result from the RFP images was converted into the traction force of the hESCs using unconstrained Fourier transform traction microscopy. The traction data were used to calculate the tensional stress within the colony using monolayer stress microscopy.

### Microarray analysis

Total RNA was extracted using TRIzol from the pooled samples of six stages including ES, SCD2, SCD3, FGFD3, FGFD6, and DMD8. RNA purity and integrity were evaluated using the OD 260/280 ratio and analyzed using the Agilent 2100 Bioanalyzer (Agilent Technologies, Palo Alto, USA). According to the manufacturer’s protocol, the Affymetrix Whole transcript Expression array process was performed (GeneChip Whole Transcript PLUS reagent Kit). First, cDNA was synthesized using the GeneChip Whole Transcript (WT) Amplification kit as described by the manufacturer. Afterward, using the GeneChip WT Terminal labeling kit, the sense cDNA was fragmented and biotin-labeled with terminal deoxynucleotidyl transferase. At 45°C, approximately 5.5 μg of labeled DNA target was hybridized to the Affymetrix GeneChip Human 2.0 ST Array for 16 h. After washing, hybridized arrays were stained on a GeneChip Fluidics Station 450 and scanned using a GCS3000 Scanner (Affymetrix). Signal values were computed using the Affymetrix® GeneChip™ Command Console software. The data were subsequently normalized using the robust multi-average (RMA) method implemented in Affymetrix® Power Tools. The result was exported with gene level RMA analysis, and the differentially expressed gene analysis was employed. The statistical significance of the expression data was determined by fold change. We focused on genes variable along cellular differentiation toward spinal cord development. Variable genes were identified by calculating the median absolute deviation (MAD) across all samples with the cutoff of the top 5% of the MAD values, which resulted in approximately 1,400 genes. These genes were then divided into two clusters, up- and down-regulated genes on differentiation, which were used for the analysis of functional enrichment in the GO terms for biological processes. For enrichment analysis, we used Enrichr’s web-based enrichment analysis tool^42, 43^ using the 250 most variable genes across samples.

### scRNA-seq analysis

Samples of organoids at FGFD1, FGFD6 and DMD10 stages were pooled and chopped into small pieces. Organoid pieces were then digested in dispase for 10 min at 37°C and dissociated into single cells. Further, the Chromium Controller was used to prepare libraries according to the 10X Single Cell 3’ v3 protocol (10x Genomics, Pleasanton, USA). After washing, cells were diluted in nuclease-free water to achieve a targeted cell count of 10,000 and mixed with a master mix. The mixture was loaded with Single Cell 3′ Gel Beads and Partitioning Oil into a Single Cell 3′ Chip. RNA transcripts from single cells were uniquely barcoded and reverse-transcribed within the droplets. Subsequently, cDNA molecules were pooled, and the cDNA pool went through an end repair process, the addition of a single ‘A’ base, and ligation of the adapters. The products were then purified and enriched with PCR to create the final cDNA library. The purified libraries were quantified using qPCR according to the qPCR Quantification Protocol Guide (KAPA) and qualified using the Agilent Technologies 4200 TapeStation (Agilent Technologies). Subsequently, the libraries were sequenced using HiSeqX (Illumina) with a read length of 28 bp for read 1 (cell barcode and UMI), 8 bp index read (sample barcode), and 91 bp for read 2 (RNA read).

Single-cell gene expression data was analyzed using Cell Ranger v5.1.0. Briefly, raw BCL files from Illumina sequencing instruments were demultiplexed to generate FASTQ files using “cellranger mkfastq.” These raw FASTQ files were then analyzed using “cellranger count.” The “cellranger count” step included mapping to the human reference genome (GRCh38), measuring gene expression with the unique molecular identifier (UMI) and cell barcode, determining cell clusters, and conducting differential gene expression analysis. The final dataset was built by aggregating multiple independent samples using “cellranger aggr.”

Raw count matrices of single cell data were imported into Seurat 4.0.2^64^. Further analysis was performed using R functions, unless otherwise noted. For quality control, we filtered out from downstream analysis (i) genes expressed in less than 10 cells, (ii) cells with over 15% reads mapping to mitochondrial genes to avoid dying cells, and (iii) cells with more than 8,000 or fewer than 200 uniquely detected genes in each cell to discard data from empty droplets or doublets. The raw count values of each sample were then log-normalized using logNormCounts::scater^65^ and then scaled using ScaleData::Seurat with default parameters.

Next, we used RunHarmony::harmony to remove variations from different runs for each sample without losing biological variations. Then we selected highly variable genes (HVGs) using getTopHVGs::scran^66^ for cell clustering. The genes with the false discovery rate (FDR) < 0.01 were identified as HVGs. Cell clustering was performed with FindCluster::Seurat using the top 20 principal components (PCs) of 1,424 HVGs, and the result was visualized using UMAP version 0.5.1.

### Cell-type annotation

We carried out cell-type annotation for the 20 clusters previously obtained. First, we tried to annotate the cluster with the expression of conventional markers following the snapshot diagram of spinal cord development by Alynick *et al*. Next, for an unbiased cell type assignment, we used Seurat v3^52^ data integration and label transfer method, following the same protocol used in our previous study^6^. Briefly, cell-type assignment was performed in a two-step process using the “Mouse Spinal Cord Atlas” dataset as the reference. The coarse level assignment identified progenitor cells and neurons, whereas the fine level analysis included all dorsal/ventral subclasses (Fig. S5). Cell types with the highest prediction score were assigned to each cluster initially. We then examined the expression of known marker genes of the assigned cell type for each cluster. For clusters with concordant gene expression pattern, we maintained the subclass assignment. Otherwise, we annotated it out at the coarse level. Default parameters and 30 PCs were used in all Seurat functions.

### Trajectory analysis

To elucidate the differentiation processes of cells in our organoid, we performed a single-cell trajectory analysis using Monocle3 (version 1.0.1) that inferred an evolutionary trajectory and placed cells at proper positions along the trajectory (pseudotime)^48^. The normalized data from Seurat was used as input to Monocle3. First, we clustered the cells using cluster_cells() function to separate partitions assuming the common transcriptional ancestry, which predicted only one partition in the dataset. Next, we used learn_graph() function to fit principal graph and then ordered the cells using order_cells() function. We chose the point with the most FGFD1 samples as the starting point.

To identify genes that determined the dorsal and ventral fates during early differentiation, we took the subset of the branches belonging to the neural progenitor cluster. We observed the differentiation of neural progenitors into mitotic dorsal and mitotic ventral axes along the trajectory in an outgoing way (from the middle to both sides). Then, we obtained genes that exhibited differential expression in the pseudotime along the trajectory using the graph_test() function. FDR < 0.01 yielded 1,617 statistically significant genes, which were further grouped into 6 clusters according to the pseudotemporal expression pattern.

### Spatial transcriptome analysis

Frozen samples were embedded in OCT compound (VWR, USA) and sectioned at −20°C with a cryotome (Thermo Scientific, USA). Tissue sections were placed on chilled Visium Tissue Optimization (1000193, 10X Genomics, USA) and Visium Spatial Gene Expression (1000184, 10X Genomics), and attached to the slides by warming on heating block. We fixed the sections onto the slides in chilled methanol for staining under the Visium User Guide (10X Genomics, USA). For gene expression analysis, cDNA libraries were prepared according to the Visium Spatial Gene Expression User Guide and sequenced on a NextSeq 550 System (Illumina, USA) at a sequencing depth up to 250 M read-pairs per sample.

Raw FASTQ files and histology images were processed using the Space Ranger software v1.0.0, which uses STAR v.2.5.1b ^67^ for genome alignment against the Cell Ranger hg38 reference genome “refdata-cellranger-GRCh38-3.0.0,” available at: http://cf.10xgenomics.com/supp/cell-exp/refdata-cellranger-GRCh38-3.0.0.tar.gz. The alignment and count processes were performed by “spaceranger count,” which commenced with specifying the input of FASTQ files, reference, section image, and Visium slide information. The pipeline detects a tissue area by aligning the image to the printed fiducial spot pattern of a Visium slide and recognizing stained spots from the image.

The Visium spatial transcriptome data were analyzed by closely following the computational pipeline for standard single-cell transcriptome data. The raw count matrix of VISIUM data was loaded into Seurat 3.2.1. We then discarded spots of low quality that contained more than 6% of their reads mapping to mitochondrial genes (dying cells) or that had less than 2,000 uniquely expressed genes (insufficient cells). The remaining 53 spots were assumed to be of good quality and used in the downstream analysis. After normalization and scaling, spots were clustered using the 20 most significant PCs as before, which yielded 2 clusters of spots. In an effort to characterize these two clusters, we performed a composite analysis that integrated the spatial transcriptomic data with scRNA-seq data as follows:

1. scRNA-seq data for the DMD10 sample that matched the spatial transcriptome data were used as a reference. Thus, we re-processed the scRNA-seq data for the DMD10 sample separately and obtained cell clusters. Annotations of cell states (mitotic or postmitotic), dorsal and ventral axis, and sections in each state were carried out in the same way as before.
2. We performed the label transfer process by following the “reference-based” integration workflow of Seurat. Briefly, we normalized each dataset with SCTransform and integrated two datasets using FindTransferAnchors. Subsequently, the discrete labels from scRNA-seq clustering and annotation were transferred to the spatial transcriptomic data using the TransferData function of Seurat.
3. The above procedure assigns a probabilistic score (between 0 and 1) of each scRNA-seq-derived label for each spot. Thus, we obtain a series of probabilistic scores representing diverse cell states and sections, which could be used to characterize each spot in the spatial transcriptome data.

We further annotated each spot according to the marker genes in the Snapshot on the spinal cord development ^46^. Each spot was assigned to a specific section if the scaled expression of a unique section marker was greater than 1.

## ACKNOWLEDGEMENTS

**Funding:** This research was supported by the National Research Foundation of Korea (NRF) grant, funded by the Korean government (MSIP) (NRF-2019M3D1A1078940, NRF-2017M3A9B3061308, NRF-2017M3C7A1047654, and NRF-2016R1D1A1B01011346). **Author contributions:** K.S. and J.R.R.: conception, design of the work, acquisition, analysis, interpretation of data, have drafted work and substantially revised it. S.C. and S.L.: analysis, interpretation of data, have drafted the work. J-H.L., B.L., Y.K., H.M.C., Y.J.K., E.Y., D.G. and H.K.: resources. J.H.K.: analysis. H.J.: acquisition, analysis, interpretation of data. Y.P.: interpretation of data. H.Y.K., T.L. and W.-Y.P.: acquisition. W.S.: conception, design of the work, interpretation of data, have drafted the work and substantially revised it. **Declaration of interest:** The authors declare no competing interests.

