## supplementary figures and tables for "Symmetry breaking of hPSCs in micropattern generates a polarized spinal cord-like organoid (pSCO) with dorsoventral organization"

### Suppl Figure 1-Figure 1 related

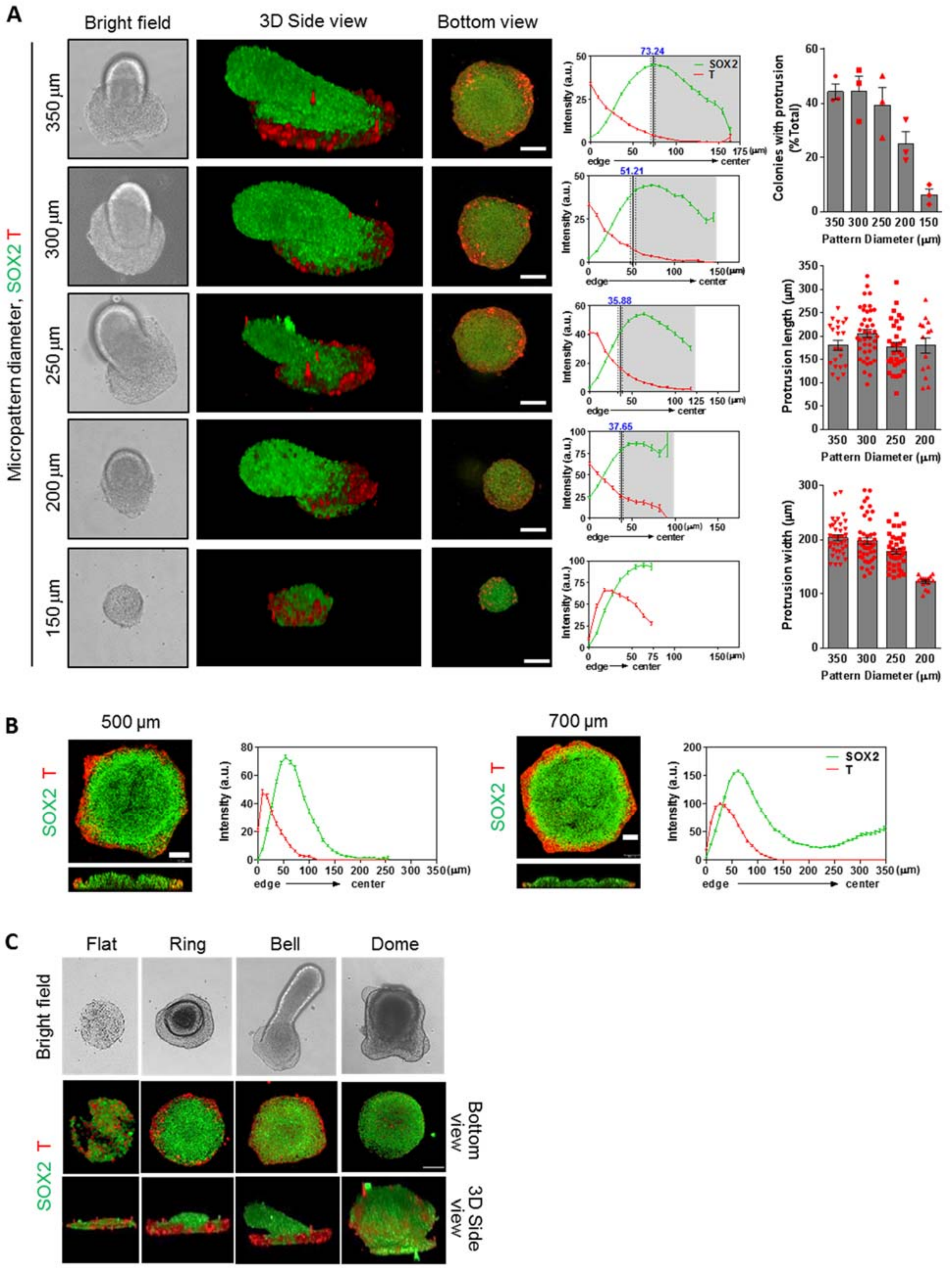


Figure S1. Geometric factors affecting cell specification and morphogenesis in micropatterned colonies treated with SB/Chir. Effects of micropattern diameter between 150 and 350 μm (A), and 500 and 700 μm (B) on spatial cell patterning and colony morphogenesis. Immunofluorescence analysis and quantification of colonies grown on micropatterns with the indicated diameters at Day 3 with SB/Chir treatment. Confocal images were taken in 5-μm steps along the z-axis after fixation and immunostaining with the indicated antibodies, and three-dimensional renderings were created. Quantification of fluorescent intensities at each position shown as mean ± SEM. Numbers in blue color at the top of each intensity graphs indicate the distance from the boundary of the position at which colony protrusion occurred in the micropatterned colony. Percentage of colonies with the protrusion, and length and width of the protrusion were quantified. (C) Spatial cell patterning and colony morphogenesis in micropatterned colonies. Immunofluorescence analysis of colonies grown on micropatterns at Day 3 with SB/Chir treatment. Confocal images were taken in 5-μm steps along the z-axis after fixation and immunostaining with the indicated antibodies, and three-dimensional renderings were created. All images are representative examples from at least three independent experiments. Scale bar, 100 μm.

#### Suppl Figure 2-Figure 1 related

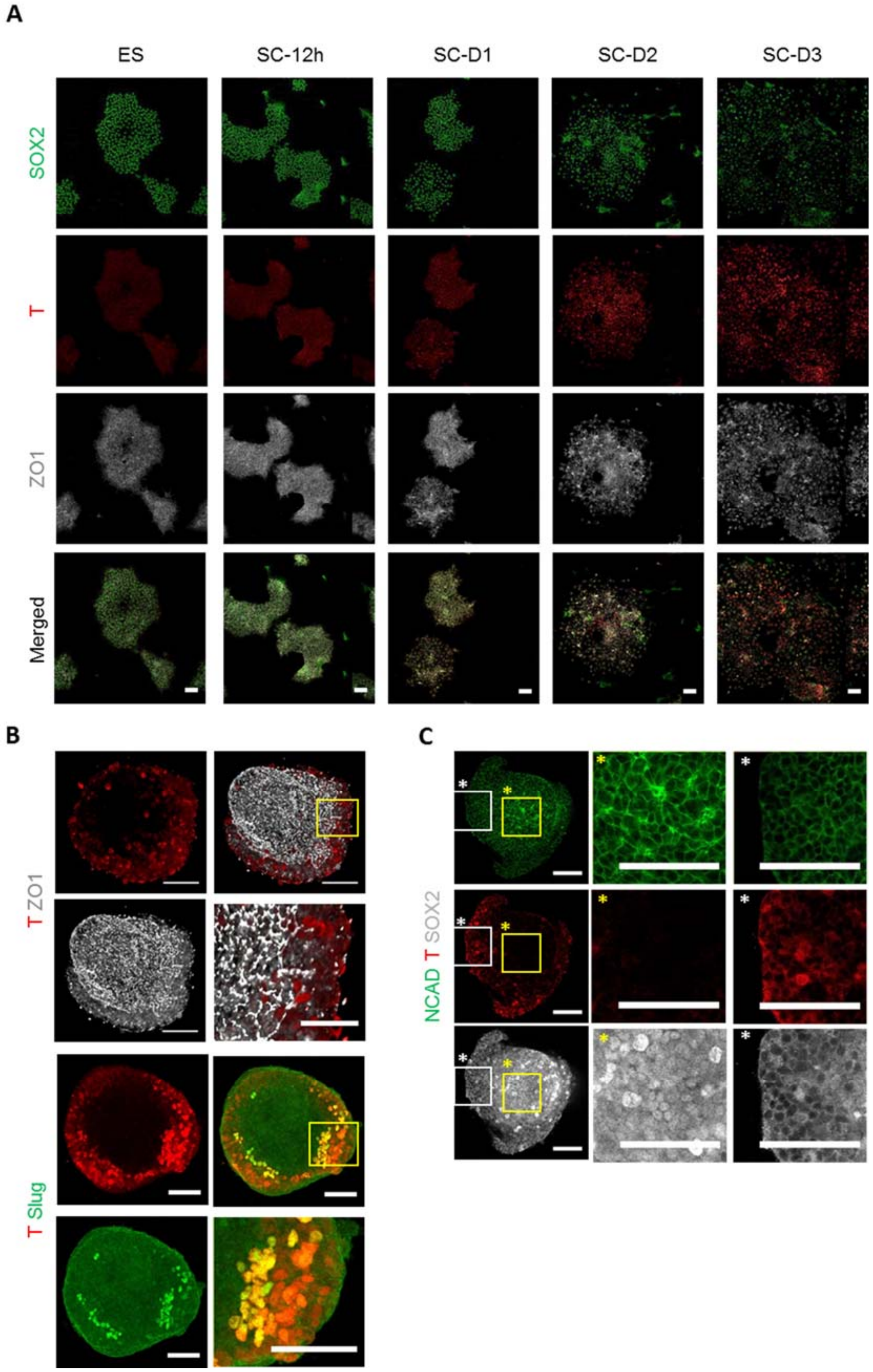


Figure S2. Characterization of differentiated hESCs. (A) Time-course immunofluorescence analysis of monolayer hESC differentiation upon SB/Chir treatment under standard culture. Confocal images were taken after fixation and immunostaining with the indicated antibodies. Scale bar, 100 μm. (B) Loss of tight junction in micropatterned colonies treated with SB/Chir at Day 3. Confocal images were taken in 1- or 5-μm steps along the z-axis after fixation and immunostaining with the indicated antibodies, and stacked using Z-stack maximum projection. High-magnification images correspond to inset. (C) NCAD expression in micropatterned colonies treated with SB/Chir at Day 3. Confocal images were taken after fixation and immunostaining. High-magnification images correspond to inset. All images are representative examples from at least three independent experiments. Scale bar, 100 μm.

#### Suppl Figure 3-Figure 2 related

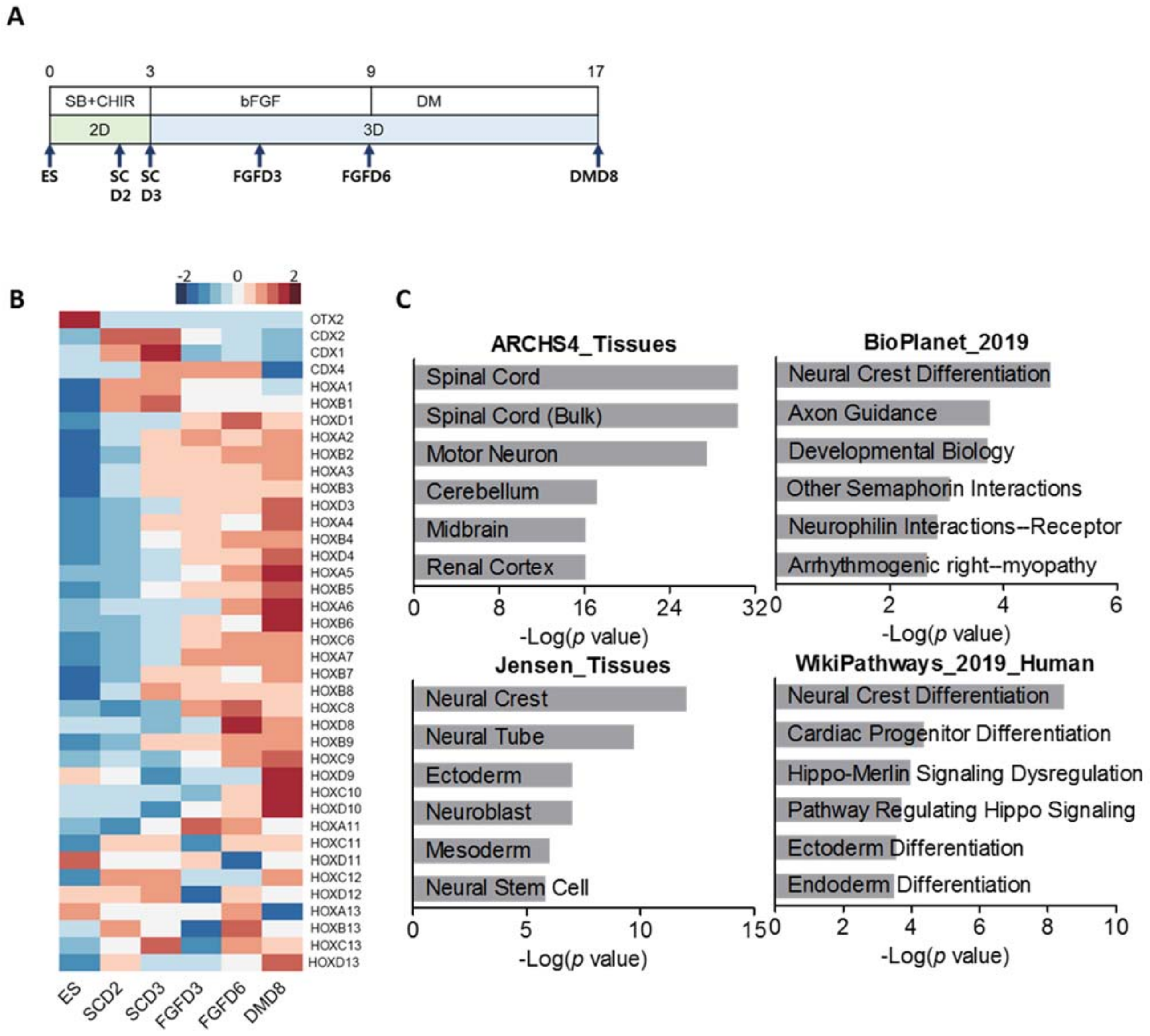


Figure S3. Microarray analysis. (A) Schematic of the time points at which samples for microarray analysis were collected. Pooled samples at each group were used for microarray analysis. (B) Heatmap of HOX gene expression in organoids over time. (C) Enrichr analysis of Top 250 most variable genes.

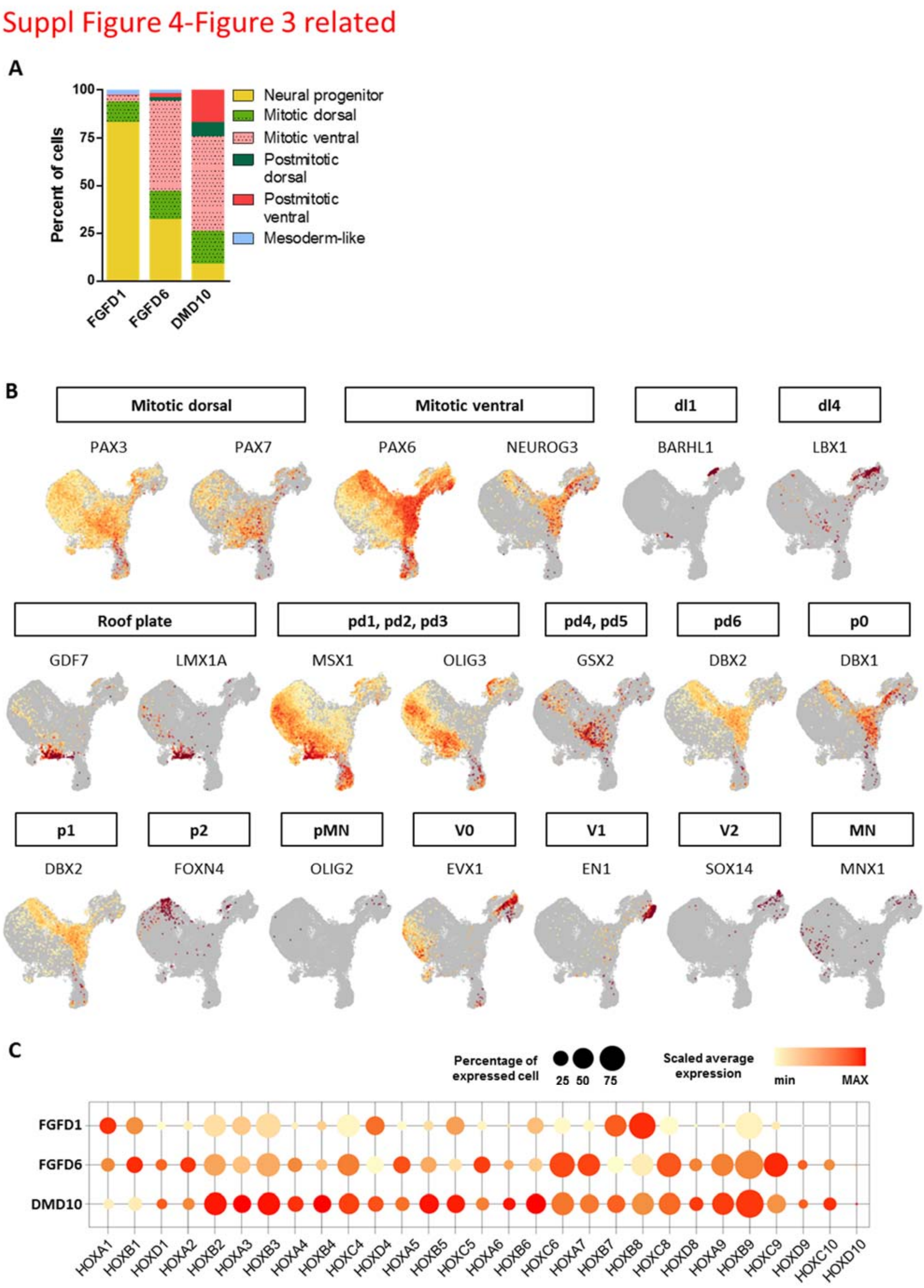


Figure S4. Single cell RNA-sequencing analysis. (A) Percentage of cells of each cluster in FGFD1, FGFD6 and DMD10 cells are differently colored. (B) Feature plots colored on the basis of gene expression of representative marker genes used to assign cluster identities. (C) Dot plots of posterior specific HOX genes across samples. The size of each circle indicates the percentage of cells where the indicated gene was detected, and the color intensity reflects the scaled average expression level within cells.

Suppl Figure 5-Figure 3 related

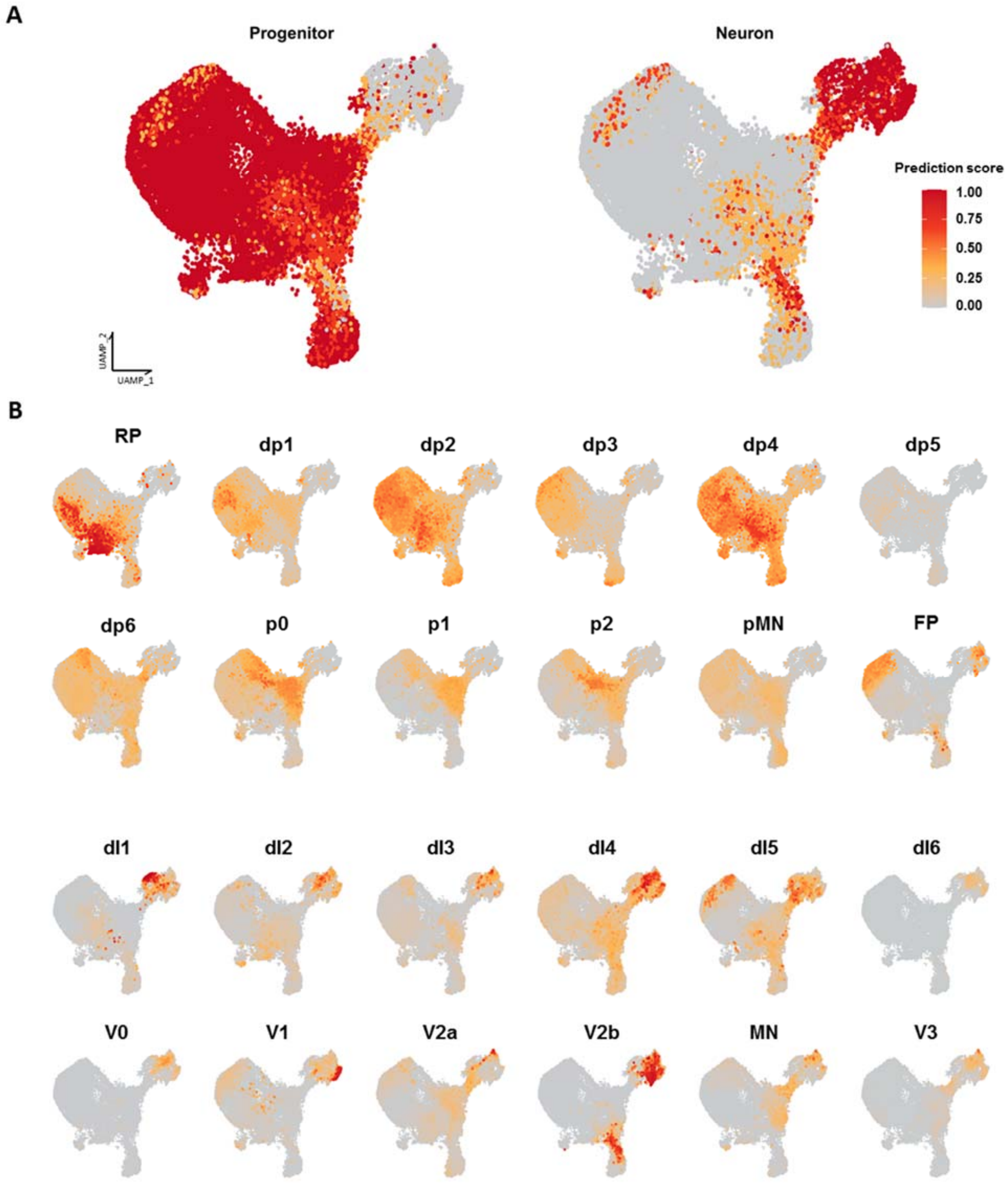


Figure S5. Comparison of pSCOs to developing mouse spinal cord. UMAP plots shows the cell assignment by prediction score analysis from scRNA sequencing clusters from previously published developing mouse spinal cord. Progenitor and neuron clusters (A) and detailed mitotic and postmitotic SC clusters (B).

### Suppl Figure 6-Figure 4 related

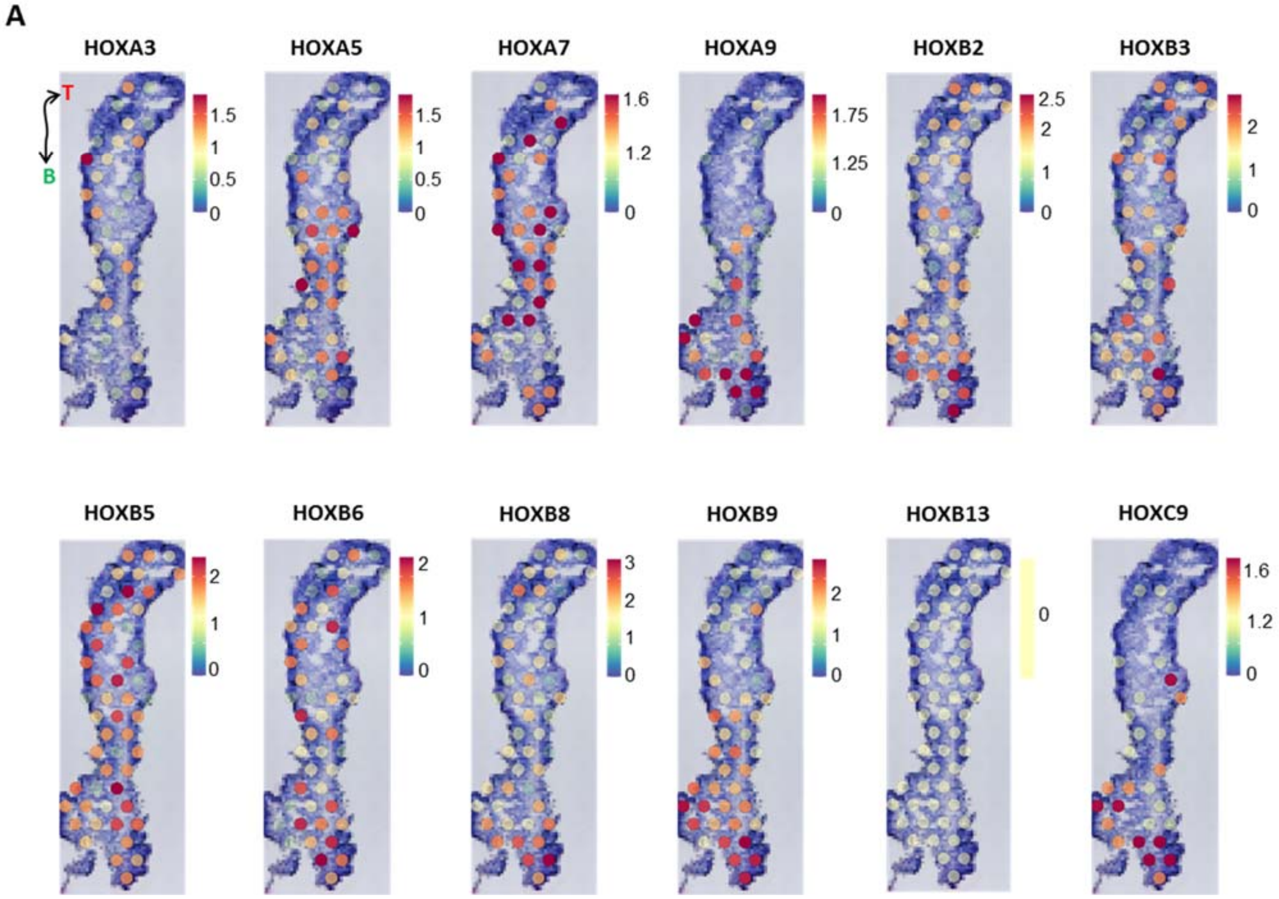


Figure S6. Analysis of HOX gene expression pattern. (A) Mapping of HOX genes in the organoid section. Black double arrowhead is labeled as T and B indicates the long axis of organoid section.

#### Suppl Figure 7-Figure 6 related

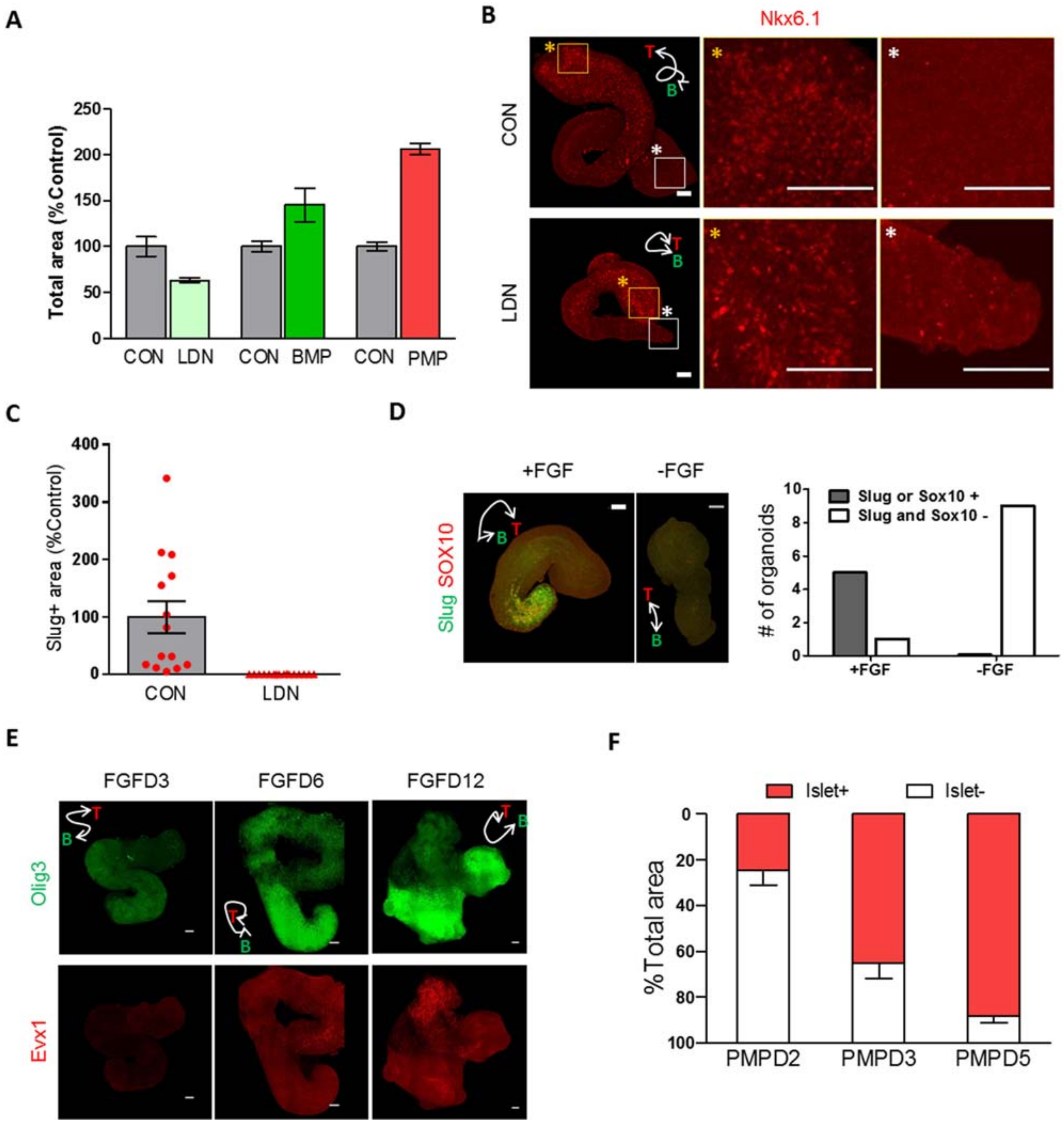


Figure S7. Effect of BMP and Shh signaling activities on DV patterning of organoids. (A) Effect of BMP inhibitor (LDN), BMP, and Shh activator (PMP) on the growth of organoids. Quantification of the total area in each group shown as mean ± SEM. (B) Effect of BMP inhibitor (LDN) on the Nkx6.1-positive region and the Nkx6.l expression level. Confocal images were taken in 5-μm steps along the z-axis after fixation and immunostaining with indicated antibodies, and stacked using Z-stack maximum projection. High-resolution images correspond to insets. White double arrowhead labeled as T (top side) and B (bottom side) indicates the long axis of organoids. Scale bar, 100 μm. (C) Effect of BMP inhibitor (LDN) on slug-positive area in the organoid. Confocal images were taken in 5-μm steps along the z-axis after fixation and immunostaining with the indicated antibodies, and stacked using Z-stack maximum projection. Quantification of the slug-positive area shown as mean ± SEM. (D) Effect of bFGF on the generation of neural crest cells in organoids. Organoids were grown in the presence or absence of bFGF for 6 days. Confocal images were taken in 5-μm steps along the z-axis after fixation and immunostaining with the indicated antibodies, and stacked using Z-stack maximum projection. Numbers of organoids with or without Slug- or SOX10-positive regions were quantified. Scale bar, 100 μm. (E) Effect of the duration of bFGF treatment on DV axis organization in organoids. Organoids were immunostained with the dorsal (Olig3) and ventral (Evx1) markers. Confocal images were taken in 5-μm steps along the z-axis after fixation and immunostaining with the indicated antibodies, and stacked using Z-stack maximum projection. Scale bar, 100 μm. (F) Effect of the duration of Shh activator (PMP) treatment on Islet-positive region in organoids. After PMP treatment for the indicated times, organoids were immunostained with anti-Islet1/2 antibody. Confocal images were taken in 5-μm steps along the z-axis and stacked using Z-stack maximum projection, and the Islet-positive area was quantified. Quantification of islet-positive area in the organoid (% total area) shown as mean ± SEM. All images are representative examples from at least three independent experiments.

Movie S1. Emergence of center protrusion at the colony during the 3rd day of SB/Chir treatment

Movie S2. Time-course changes in colony morphology and spatial patterning of SOX2- and T-positive cells during SB/Chir treatment

Movie S3. Four types of colony morphology immunostained with anti-SOX2 and anti-T antibodies at day 3 of SB/Chir treatment

Movie S4. Effect of micropattern diameters on colony morphology and spatial patterning of SOX2- and T-positive cells at day 3 of SB/Chir treatment

Movie S5. Growth of 3D structures after detachment from 2D micropatterned substrate

Movie S6. Rosette structures immunostained with anti-ZO1 antibody at SC-D3, FGFD1, and FGFD6

Movie S7. Dorsal and ventral patterning of spinal cord organoids immunostained with markers for spinal cord progenitor cells

Movie S8. Dorsal and ventral patterning of spinal cord organoids immunostained with markers for spinal cord postmitotic neurons

**Table S1. Key resources**

| REAGENT or RESOURCE | SOURCE | IDENTIFIER |
|---|---|---|
| Antibodies | | |
| Rabbit polyclonal anti-Sox2 | Millipore | Cat# AB5603; RRID: AB_2286686 |
| Mouse monoclonal anti-Sox2 | Santa Cruz Biotechnology | Cat# sc-365823; RRID: AB_10842165 |
| Goat polyclonal anti-Brachyury | R&D Systems | Cat# AF2085; RRID: AB_2200235 |
| Rabbit polyclonal anti-beta-Catenin | Sigma-Aldrich | Cat# C2206; RRID: AB_476831 |
| Rabbit polyclonal anti-ZO-1 | Thermo Fisher Scientific | Cat# 61-7300; RRID: AB_2533938 |
| Rabbit monoclonal anti-CDX2 | Abcam | Cat# ab76541; RRID: AB_1523334 |
| Goat polyclonal anti-Doublecortin(DCX) | Santa Cruz Biotechnology | Cat# sc-8066; RRID: AB_2088494 |
| Goat polyclonal anti-Nkx6.1 (N-15) | Santa Cruz Biotechnology | Cat# sc-15027; RRID: AB_650286 |
| Mouse monoclonal anti-Pax3 | R&D Systems | Cat# 274212; RRID: AB_2159398 |
| Mouse monoclonal anti-Pax6 | DSHB | Cat# pax6; RRID: AB_528427 |
| Rabbit monoclonal anti-Olig3 | Abcam | Cat# Ab129197; RRID: AB_11142825 |
| Mouse monoclonal anti-Pax7 | DSHB | Cat# pax7; RRID: AB_528428 |
| Rabbit monoclonal anti-Slug (C19G7) | Cell Signaling Technology | Cat# 9585; RRID: AB_2239535 |
| Mouse monoclonal anti-Sox10 (A-2) | Santa Cruz Biotechnology | Cat# sc-365692; RRID: AB_10844002 |
| Mouse monoclonal anti-Evx1/2 | DSHB | Cat# 99.1-3a2; RRID: AB_528231 |
| Goat polyclonal anti-Brn3a (C-20) | Santa Cruz Biotechnology | Cat# sc-31984; RRID: AB_2167511 |
| Rabbit polyclonal anti-Islet1 | Abcam | Cat# Ab20670; RRID: AB_881306 |
| Sheep polyclonal anti-Chx10 (N-terminus) | Millipore | Cat# AB9016; RRID: AB_2216009 |
| Goat polyclonal anti-Chat | Millipore | Cat# AB144P; RRID: AB_2079751 |
| Mouse monoclonal anti-Lim 1+2 / LhxV5 | DSHB | Cat# 4F2; RRID: AB_531784 |
| Rabbit polyclonal anti-Peripherin | Abcam | Cat# ab4666; RRID: AB_449340 |
| Mouse monoclonal anti-N-Cadherin | BD Biosciences | Cat# 610921; RRID: AB_398236 |
| Hoechst33342 | Invitrogen | Cat# H3570; CAS: 23491-52-3 |
| Donkey anti-Mouse IgG (H+L) Antibody, Alexa Fluor 488 | Thermo Fisher Scientific | Cat# A21202; RRID: AB_141607 |
| Donkey anti-Rabbit IgG (H+L) Antibody, Alexa Fluor 488 | Thermo Fisher Scientific | Cat# A21206; RRID: AB_2535792 |

| | | |
|---|---|---|
| Donkey anti-Goat IgG (H+L) Antibody, Alexa Fluor 488 | Thermo Fisher Scientific | Cat# A11055; RRID: AB_2534102 |
| Donkey anti-Sheep IgG (H+L) Antibody, Alexa Fluor 488 | Thermo Fisher Scientific | Cat# A21448; RRID: AB_2534082 |
| Donkey anti-Mouse IgG (H+L) antibody, Cy3 | Jackson ImmunoResearch | Cat# 715-165-151; RRID: AB_2315777 |
| Donkey anti-Rabbit IgG (H+L) antibody, Cy3 | Jackson ImmunoResearch | Cat# 711-165-152; RRID: AB_2307443 |
| Donkey anti-Goat IgG (H+L) antibody, Cy3 | Jackson ImmunoResearch | Cat# 705-165-147; RRID: AB_2307351 |
| Donkey anti-Mouse IgG (H+L) antibody, Alexa Fluor 647 | Jackson ImmunoResearch | Cat# 715-606-150; RRID: AB_2340865 |
| Donkey anti-Rabbit IgG (H+L) antibody, Alexa Fluor 647 | Jackson ImmunoResearch | Cat# 711-605-152; RRID: AB_2492288 |
| Donkey anti-Goat IgG (H+L) antibody, Alexa Fluor 647 | Thermo Fisher Scientific | Cat# A21447; RRID: AB_2535864 |
| Chemicals, Peptides, and Recombinant Proteins | | |
| SB 431542 | TOCRIS | Cat# 1614; CAS: 301836-41-9 |
| CHIR 99021 | Sigma-Aldrich | Cat# SML1046; CAS: 252917-06-9 |
| LDN-193189 | Stemgent | Cat# 04-0074; CAS: 1062368-24-4 |
| Human bFGF | R&D Systems | Cat# 233-FB; GenPept: P09038 |
| Human BMP2 | PEPROTECH | Cat# 120-02; GenPept P12643 |
| Purmorphamine | Sigma-Aldrich | Cat# 540220; CAS: 483367-10-8 |
| Matrigel hESC-Qualified Matrix | Corning | Cat# 354277 |
| Cell Recovery Solution | Corning | Cat# 354253 |
| mTeSR1 | STEMCELL Technologies | Cat# 85850 |
| ReLeSR | STEMCELL Technologies | Cat# 05872 |
| DMEM/F-12 | Gibco | Cat# 11320033 |
| N2 | Gibco | Cat# 17502048 |
| B27 | Gibco | Cat# 17504044 |
| Nonessential amino acids (NEAA) | Gibco | Cat# 11140050 |
| penicillin/streptomycin (P/S) | Gibco | Cat# 15140122 |
| β-mercaptoethanol | Gibco | Cat# 21985023 |
| HBSS | Gibco | Cat# 14175095 |
| Experimental Models: Cell Lines | | |
| Human: Passage 30 H9 ES cells | WiCell | WA09; WAe009-A |
| Software and Algorithms | | |
| ImageJ-Fiji | Schneider et al., 2012 | https://imagej.net/Fiji |
| LasX | Leica | https://www.leica-microsystems.com/products/microscope-software/p/leica-las-x-ls/ |

| | | |
|---|---|---|
| Prism5 | GraphPad | https://www.graphpad.com/ |
| MATLAB | MathWorks | https://kr.mathworks.com/products/matlab.html |
| SigmaPlot 12 | SYSTAT | https://systatsoftware.com/downloads/download-sigmaplot/ |
